# Mathematical modeling of ventilator-induced lung inflammation

**DOI:** 10.1101/2020.06.03.132258

**Authors:** Sarah Minucci, Rebecca L. Heise, Michael S. Valentine, Franck J. Kamga Gninzeko, Angela M. Reynolds

## Abstract

Respiratory infections, such as the novel coronavirus (SARS-COV-2) and other lung injuries, damage the pulmonary epithelium. In the most severe cases this leads to acute respiratory distress syndrome (ARDS). Due to respiratory failure associated with ARDS, the clinical intervention is the use of mechanical ventilation. Despite the benefits of mechanical ventilators, prolonged or misuse of these ventilators may lead to ventilation-associated/ventilation-induced lung injury (VILI). Damage caused to epithelial cells within the alveoli can lead to various types of complications and increased mortality rates. A key component of the immune response is recruitment of macrophages, immune cells that differentiate into phenotypes with unique pro- and/or anti-inflammatory roles based on the surrounding environment. An imbalance in pro- and anti-inflammatory responses can have deleterious effects on the individual’s health. To gain a greater understanding of the mechanisms of the immune response to VILI and post-ventilation outcomes, we develop a mathematical model of interactions between the immune system and site of damage while accounting for macrophage polarization. Through Latin hypercube sampling we generate a virtual cohort of patients with biologically feasible dynamics. We use a variety of methods to analyze the results, including a random forest decision tree algorithm and parameter sensitivity with eFAST. Analysis shows that parameters and properties of transients related to epithelial repair and M1 activation and de-activation best predicted outcome. Using this new information, we hypothesize interventions and use these treatment strategies to modulate damage in select virtual cases.

## 1. Introduction

Inflammation occurs in the lungs when an immune response is initiated to eliminate an insult. Types of insults include inhaled pathogens, such pneumonia, tuberculosis, SARS-COV-2, or other harmful particles. In the most severe cases this leads to acute respiratory distress syndrome (ARDS). Due to respiratory failure associated with ARDS, the clinical intervention is the use of mechanical ventilation. When individuals have a severe form of COVID-19, the disease caused by SARS-COV-2, the disease can lead to respiratory failure and death of the patients. In a recent study, two-thirds of patients admitted for COVID-19 required mechanical ventilation [1].

Despite the benefits of mechanical ventilators, prolonged or misuse of these ventilators may lead to ventilation-induced lung injury (VILI). In this work we will focus on the tissue damage associated with mechanical ventilation and resulting immune cell recruitment. The damage caused to alveolar sacs (clusters of alveolar cells) during mechanical ventilation can lead to volutrauma (extreme stress/strain), barotrauma (air leaks), atelectrauma (repeated opening and closing of alveoli), and biotrauma (general severe inflammatory response). If the trauma increases, it can lead to multi-system organ failure [2, 3].

It has also been shown that the inflammatory response of the elderly is altered in the lungs and other areas [4, 5]. As compared to younger individuals, increased levels of circulating inflammatory cytokines and different immune cell function have been reported in older patients [6]. A 2003-2008 study conducted at Bridgeport Hospital reported that 4,238 out of 9,912 (42.8%) patients received mechanical ventilation for a median of two days. Mortality or discharge to extended-care facilities increased for each decade of age greater than 65 years [7]. Additionally, the case fatality rate for COVID-19 patients over 70 years old and over 80 years old was around 50.8% and 14.8% of the total number of deaths, respectively [8]. This is in agreement with other studies reporting higher rates of severe outcomes in patients with COVID-19 aged 65 or older [9]. The change in the inflammatory response with patient age combined with the increased need for ventilation and increased mortality rate among the elderly stresses the need to investigate the influence of aging in VILI. The framework we have built here addresses VILI with various parameters and initial conditions that can be narrowed in future studies with data from different age groups and/or insults to explore dynamics and driving factors in various diseases related to age and/or outcome.

We used mathematical modeling to investigate the role of the pulmonary immune response and treatments in ventilator-induced damage. We adapted a model developed by Torres *et al.* for the innate immune response to bacteria, which accounts for macrophage polarization, by including epithelial dynamics and stretch-induced recruitment of immune cells [10]. We use this model to understand the mechanisms by which the immune system responds to damaged epithelial cells and the sensitivity of post-ventilation outcome to components of this complex process. We begin this study by analyzing the epithelial subsystem mathematically. This allows us to understand fixed point stability and how various parameters affect stability for the new portion of the model. The full model is a large system of ordinary differential equations with a large number of parameters and a variety of nonlinear dynamics. Allowing the parameters in the model to vary over biologically feasible ranges using Latin hypercube sampling simulates the variety of immune system dynamics that may be observed in patients. We organize disease progressions into three categories, healthy, moderate inflammation, and severe inflammation, based on the percentage of healthy epithelial cells. To determine what is driving differences in outcome, we use a variety of methods to analyze the resulting dynamics: 1) comparison of parameters associated with different outcomes, 2) random forest decision tree algorithm, which parses through the variety of predictors that may be particularly important in the immune response to VILI and 3) parameter sensitivity with eFAST, a variance-based method.

### 1.1. Biological background

The alveolar epithelium consists of alveolar type I and type II cells. Alveolar type I cells make up about 95% of the alveolar surface and are primarily responsible for facilitating gas exchange. Type II cells cover the other 5% of the surface and are important in the innate immune response. In the presence of damage, these cells proliferate to repair the epithelium and can also differentiate to type I cells [11, 12]. The extent to which the alveolar epithelium is damaged is a useful indicator of the overall effects of a lung insult [13].

The immune response is divided into innate (non-specific) and adaptive (acquired) responses. Two of the most important innate immune cells are neutrophils and macrophages, which can be tissue-specific or recruited to the site upon damage. The innate response is always present and ready to defend against pathogens or other insults. On the other hand, the adaptive immune response includes B and T cells, which differentiate in such a way that they are effective at fighting specific pathogens. They are recruited by antigen-presenting cells, such as dendritic cells and macrophages, that are a part of the innate immune response.

In this work, we concentrate on the innate immune system when modeling VILI to gain a better understanding of the epithelial and immune cell interactions. Lung infection may lead to the need for mechanical ventilation and the resulting model could be adapted in the future to study mechanical ventilation with infection. Initially we consider a system in which the immune response is triggered by damage associated with the ventilator without infection.

One of the key components of this response is recruitment of macrophages from the bone marrow and bloodstream to the damaged area to support the population of resident alveolar macrophages. Macrophages send signals to other immune cells and aid in the process of eliminating dead cells and repairing damaged ones [14]. Phenotypes of macrophages can range from “pro-inflammatory” (M1) or “anti-inflammatory” (M2) based on their activators and byproducts [15, 16]. Their pro-inflammatory behavior includes destroying pathogens, consuming damaged cells, and amplification of signaling. Their anti-inflammatory response, which counteracts pro-inflammatory behavior, promotes repair by producing anti-inflammatory cytokines and removing apoptotic neutrophils. A single macrophage may produce both proinflammatory and anti-inflammatory signals concurrently, which can make classification and identification of phenotype a difficult question.

Another important type of immune cell is the neutrophil, which responds quickly to pro-inflammatory signals sent from damaged epithelial cells and other resident cells. A small amount of neutrophils are found in the lungs in homeostasis. Additional neutrophils are recruited from bone marrow in response to pro-inflammatory signals from damaged epithelial cells and resident macrophages during an insult in large numbers [17]. Neutrophils have phagocytic capabilities in the presence of invading pathogens, but in the case of VILI without infection neutrophils recruit other immune cells such as macrophages through the production of pro-inflammatory agents such as proteinases and cytokines and contribute to the removal of damaged or dead tissue. An overabundance of neutrophils and their byproducts can cause further unnecessary damage [18]. Neutrophils are relatively short-lived; they become apoptotic and are removed by macrophages [17] or become necrotic in an uncontrolled death resulting in the release of cytotoxic material [19].

An imbalance in the pro- and anti-inflammatory responses can cause complications for the individual. Furthermore, an absence of immune cells can lead to immunodeficiency and a surplus of immune cells can result in chronic inflammation [17]. Thus, it is important to understand the immune response to lung injury and the interplay between various types of cells. It is also believed that macrophages play a significant role in the impact of aging on the immune response [6, 20, 21].

### 1.2. Mathematical background

Mathematical modeling is used to capture the complexities of the immune response to epithelial cell damage, including important feedback loops and nonlinearities. Analyzing the resulting model gives insight into the driving mechanisms of this system. An *in silico* approach allows us to simulate various scenarios or new treatments, especially when *in vivo* and *in vitro* experiments to explore possible interventions to improve outcomes for patients are difficult to perform. To our knowledge, no mathematical models have described M1/M2 interactions specific to the immune response to VILI. Many models have examine the immune response to bacterial and viral infections, such as pneumonia [22–24], tuberculosis [25–27], and influenza [28–30]. Additionally, models related to smoking and asthma [31–34], mechanical ventilation [35–42], and general inflammatory stress [4, 43] have been developed, but these models generally deal with the mechanics of the airways, including airflow, pressure, and gas exchange, and how these mechanics respond to inflammation and particle inhalation without accounting for the various cells types involved in the immune response. Models have also been developed to understand and analyze the molecular mechanisms that govern the phenotype switch that macrophages undergo from pro-inflammatory to antiinflammatory, as well as other important subcellular pathways [29, 44, 45].

Common modeling approaches used in these papers include agent-based models [27, 31, 34], partial differential equations [42, 43], ordinary differential equations [22–25, 30, 32], and Boolean models [29]. Each technique has its advantages and disadvantages, but we choose to model the inflammatory response to VILI, specifically the resulting damage to epithelial cells, using a set of coupled ordinary differential equations (ODEs), which we describe further in the following section. Systems of ODEs are ideal for modeling dynamical systems because of their ability to capture, with reasonable computation times, the highly nonlinear behavior of the many immune cells, epithelial cells and other mediators involved in the immune response to VILI. This allows for mathematical and sampling approaches to be used to determine key components of the the biological process being modeled.

## 2. Methods & Model Development

### 2.1. Epithelial subsystem

The primary focus of this model is to examine the effects of damage on the alveolar epithelium, in particular alveolar type II cells, since they are responsible for restoration of the epithelium. In this section we begin with a simple model, concentrating on the novel aspect of incorporating epithelial cells and relative damage due to inflammation. We then add variables to more accurately model the dynamics within this system.

We begin with a small three-dimensional system of differential equations, shown in Eqs (1)–(3), where *E*_*h*_ is the proportion of the local space filled by healthy cells, *E*_*d*_ is the proportion of the local space filled by damaged cells, and *E*_*e*_ represents dead cells or empty “space” that can be replaced/filled with healthy cells. Each term represents a biological event explained by the brackets above the term. This first model includes only the baseline abilities of epithelial cells to proliferate and repair themselves in the presence of sustained damage. We do not explicitly model proliferating and non-proliferating cells; the parameter *p* is modulated to reflect the general mechanism by which neighboring epithelial cells renew surrounding “space” (tracked by *E*_*e*_).

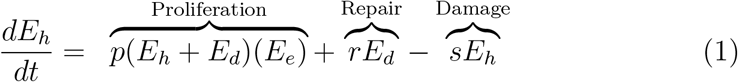

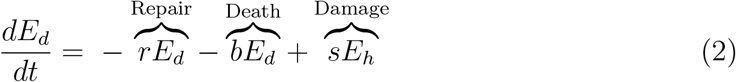

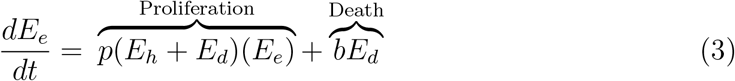

Damage from stretch due to mechanical ventilation is represented by the rate *s*, and causes healthy epithelial cells to become damaged. This general term covers over-distension for any mode of ventilation. Some damaged cells, depending on the severity of damage, have the ability to repair themselves, returning from the *E*_*d*_ state back to *E*_*h*_, represented by a baseline repair rate *r* [46]. Damaged cells may also decay naturally at a rate *b*.

The first terms in Eq (1) for *E*_*h*_, and Eq (3) for *E*_*e*_, account for proliferation of the healthy and damaged cells into empty space. Note that total local space is conserved: *E*_*e*_ + *E*_*h*_ + *E*_*d*_ = 1. Therefore, we can define *E*_*e*_ = 1 (*E*_*h*_ + *E*_*d*_) and rewrite this term, where it becomes the standard logistic growth with a carrying capacity of 1, associated with 100% of space being filled. Eliminating *E*_*e*_ gives rise to a two-dimensional system, Eqs (4)–(5).

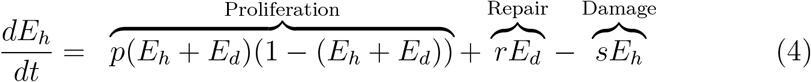

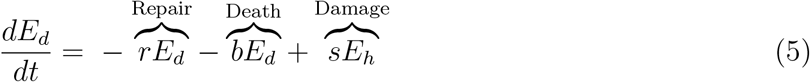

Nearby epithelial cells and progenitor cells, stem cells that can differentiate into specific types of epithelial cells only, perform this task. These cells spread and replicate to fill the empty space left by dead epithelial cells [46–48]. In this model we do not account for the progenitor cells. Therefore, we only account for proliferation associated with local epithelial cells.

Stability analysis reveals that in the absence of stretch (*s* = 0) and with all positive parameters, (0, 0) is a saddle node and (0, 1) is a stable equilibrium with eigenvalues *λ*_1_ = −*r* − *b* and *λ*_2_ = −*p*. Given a nonzero initial condition for damaged cells the epithelial cells subsystem will resolve to the fully repaired fixed point (0, 1).

In the presence of sustained stretch (*s* > 0), the *E*_*d*_ nullcline switches from a vertical line to a line with slope (*r* + *b*)*/s*. The second equilibrium point changes from (0, 1) to

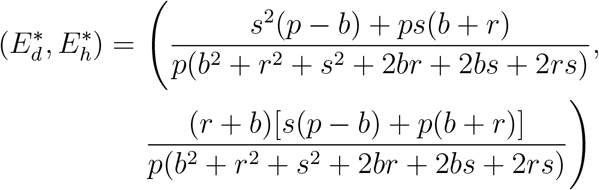

Therefore in the presence of damage, there no longer exists an equilibrium associated with full recovery.

Exploratory simulations demonstrate that there is a bifurcation with respect to *p*, the proliferation rate of epithelial cells. A bifurcation diagram for this parameter, shown in Fig 1, has one transcritical bifurcation at *p*^*^ = 0.497. The bifurcation diagrams in this manuscript were created using XPPAUT [49] with code included in the supplementary materials. In this figure, we show the proportion of space occupied by healthy epithelial cells as a percentage, which is *E*_*h*_ multiplied by 100. The second equilibrium for values of *p* below the bifurcation is not included in the diagram, since it is non-biological (negative *E*_*h*_). For small values of *p*, the ability of healthy cells to proliferate and replace dead cells is insufficient and damage causes both healthy and damaged cells to approach 0%. On the other hand, for values of *p* larger than *p*^*^, the system approaches the stable nonzero equilibrium 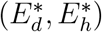, which is closer to (0, 1) for higher values of *p* even in the presence of sustained damage.

**Figure 1:**
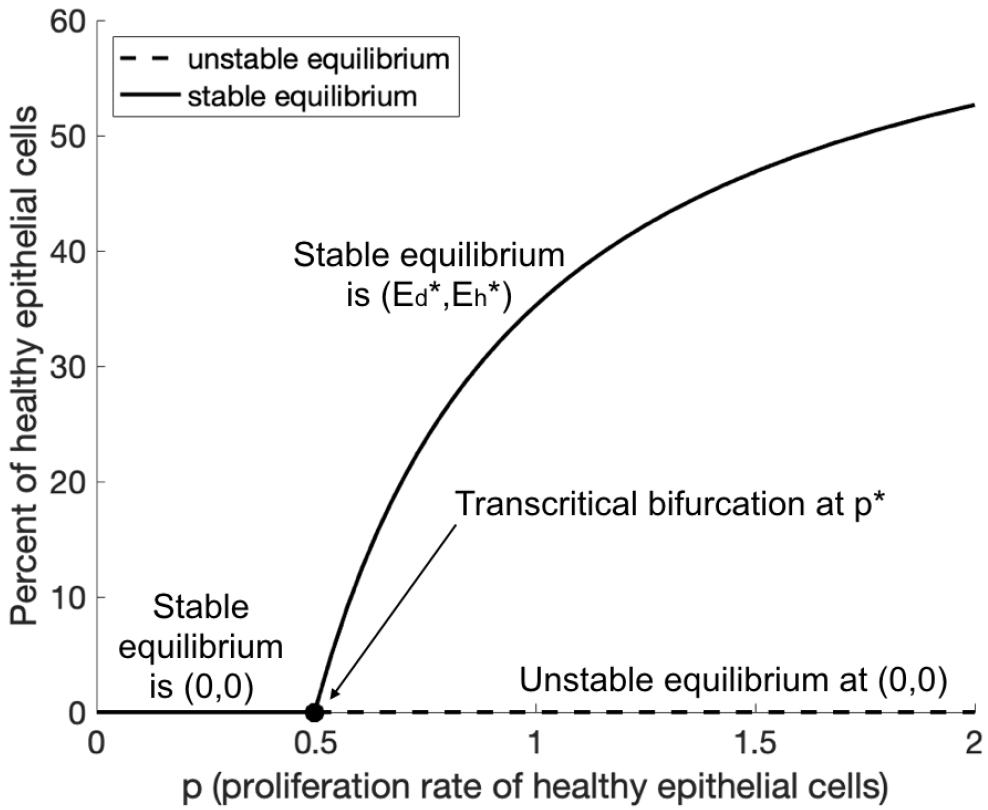
The epithelial subsystem generates a transcritical bifurcation for the parameter *p*. Bifurcation diagram for the proliferation parameter *p* for the epithelial system with stretch and no immune response. Other parameters are set to *r* = 2.6, *s* = 0.22, and *b* = 0.74. The unstable equilibrium below *p* < *p*^*^ = 0.497 is not included in the figure, since it is not biologically relevant.

### 2.2. Fixed immune response

Next we examine the roles of immune cells, especially neutrophils and macrophages, by adding several terms to Eqs (1) and (2). We first focused on dynamics with a fixed immune response, because when we work with the full model (described in the next section), we only consider parameter sets that give rise to steady-state solutions in the absence of ventilator-induced damage. Therefore, we decided to start our model development by analyzing *E*_*h*_ and *E*_*d*_ with immune cells as parameters before including their full dynamics. The modifications are shown in Eqs (6) and (7).

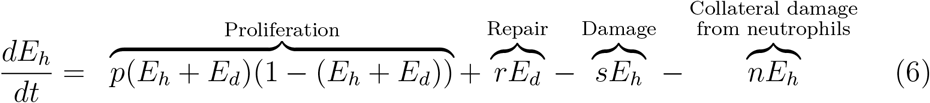

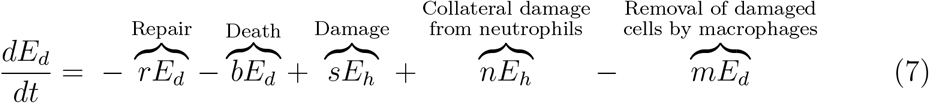

The physical presence of immune cells, especially first-responder neutrophils, causes small-scale collateral damage as they clear debris [50] and can be especially deleterious if the response is overzealous [18]. This biological event is modeled as the last term in Eq (6) with cells switching from a healthy to a damaged state at the rate *n*. M1 macrophages aid in the clearance of damaged cells to make room for replacement by new, healthy cells through subcellular signalling and phagocytosis [14, 47]. The last term in Eq (7) represents this loss of damaged cells.

The stability analysis is similar to that from the model without the immune response, with additional parameters *m, n* that can shift steepness of the nullcline or the speed at which the system approaches or diverges from an equilibrium. The parameter *p* once again plays an important role in the stability of the two critical points, (0, 0) and

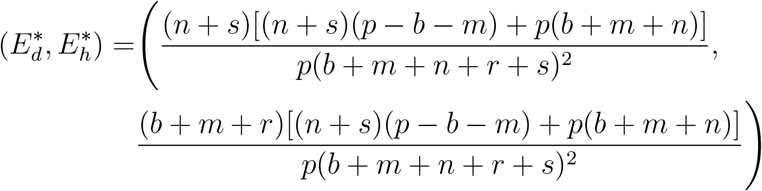

There is a transcritical bifurcation when the value of *p* is varied; given its similarly to Fig 1, it is not shown here. For the same parameter values as in Fig 1 (*r* = 2.6, *s* = 0.22, *b* = 0.74) with *m* = 0.92 and *n* = 1.6 added, we obtain the same *p*^*^ = 0.497. The main difference between these models is that the transcritical bifurcation point *p*^*^ may be lower because of the damage resulting from macrophages and neutrophils, represented by *m* and *n*. The rate of proliferation of healthy cells may need to be higher to counteract these effects.

The bifurcation diagram for scaled *E*_*h*_ versus *n* also has a transcritical bifurcation (see Fig 2a). For sufficiently low values of *n*, the nonzero critical point is stable, but for values above *n*^*^ = 1.364, (0, 0) is the stable equilibrium. Additionally, the two-parameter stability diagram shows a curve which separates the *p/n*-space into two stability regimes (see Fig 2b). For high enough values of *n* and low enough values of *p*, the system goes to zero for both variables. Biologically, this corresponds to a situation in which the ability of epithelial cells to proliferate is low and there are high levels of immune cells. On the other hand, with low levels of immune cells and a higher proliferation rate, the system limits to the nonzero equilibrium. It should be noted that for a large enough *p*, it would take an extremely high value of *n* to overpower proliferation and make (0, 0) the stable critical point. In the full system the initial conditions for our simulations will have similar properties to the type of steady state in the non-zero stable equilibrium region of Fig 2b. Varying levels of baseline inflammation exist given differences in patients’ age and past medical history.

**Figure 2:**
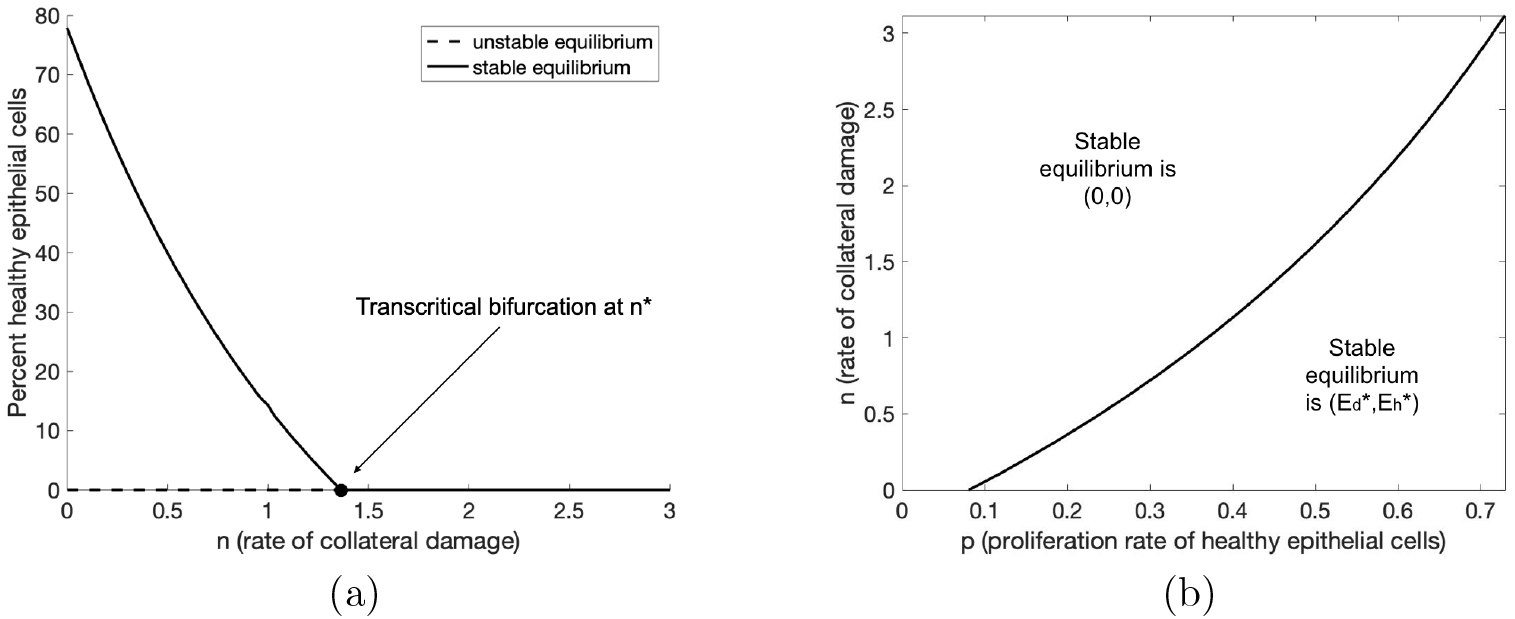
Variations on the epithelial subsystem reveal a transcritical bifurcation and two-parameter bifurcation. (a) Bifurcation diagram for epithelial subsystem when varying *n*. Other parameter values are set to *r* = 2.6, *p* = 0.45, *s* = 0.22, *b* = 0.74, *n* = 1.6, *m* = 0.92. (b) Two-parameter plot showing values of *p* and *n* which cause the subsystem to have either a zero or nonzero stable equilibrium.

These simple models provide a framework for the dynamics of the epithelium in response to damage and an introductory look into the influence of the immune response. However, there are many more complex, nonlinear interactions and events involved in VILI which we will explore in the next section.

### 2.3. Development of complete model

By adding variables to the two-dimensional system proposed above, we developed a system of coupled ordinary differential equations to model the interactions between immune cells, epithelial cells, and other mediators, shown in Fig 3. We also utilize a two-compartment method in which resident immune cells respond to the damaged epithelial cells and nonresident immune cells are recruited from the bloodstream.

**Figure 3:**
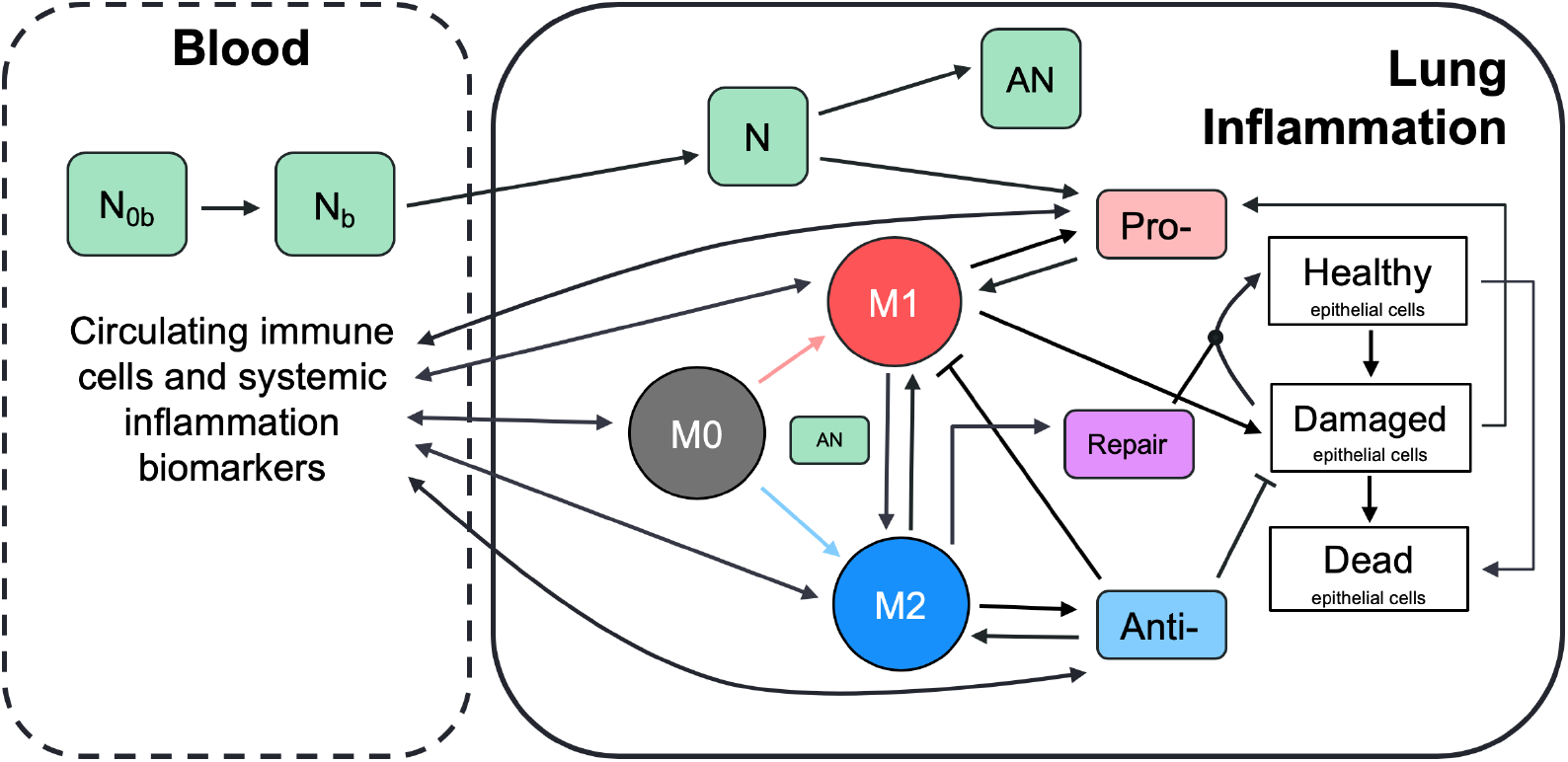
Schematic describes interactions between immune system components. Immune system components shown in this schematic are macrophages, neutrophils, various pro- and anti-inflammatory mediators, and epithelial cells. Green boxes represent various types of neutrophils, colored circles represent naive, M0, M1, and M2 macrophages. White boxes represent healthy, damaged, and dead epithelial cells/empty space (*E*_*h*_, *E*_*d*_, and *E*_*e*_, respectively). Other colored boxes represent various types of mediators that are produced by epithelial and/or immune cells and signal to immune cells. Unactivated immune cells become activated by various mediators (*p*_*b*_, *a*_*b*_ in the blood and *p*, *a* locally) and perform either pro-inflammatory or anti-inflammatory roles which are meant to remove debris (*E*_*e*_) and promote repair of damaged epithelial cells. Dynamics between cells and mediators in the blood (not shown) are similar to the detailed dynamics shown for local inflammation.

A system of ODEs is ideal for modeling these interactions because of its ability to capture distinct nonlinearities and feedback loops with relatively low computational requirements. However, one of the drawbacks of an ODE model is that it assumes a well-mixed environment, in which all elements of the model are evenly distributed throughout the given space. Biologically, this is not always the case. One way to include aspects of the spatial heterogeneity without explicitly modeling space is to use a compartmental model. Each compartment represents a well mixed environment and, when biologically appropriate, variables can move between compartments. An equation is developed for the component in each compartment in which it can be located.

Here we choose to model two compartments. The first is the site of inflammation in the lungs, specifically the epithelial cells which provide a barrier lining the alveolar cells. The second compartment is the adjacent blood vessel that provides additional immune support to the site of damage. Differentiating between these two compartments allows us to determine the concentrations of various immune cells and other mediators in each separate area and examine their movement across compartments. A two-compartmental model accounts for some spatial dynamics that a traditional system of ODEs cannot, making the model more realistic for a better understanding of the immune response to VILI.

Fig 3 gives a detailed breakdown of the dynamics in the lung. The dynamics are similar for those cells and mediators in the blood. Cell types that are tracked in each compartment are stated in Table 1. In the following subsections, we develop the equations for these variables. The parameters used in the equations are given in Table 2 with their description and range used during parameter sampling.

**Table 1:** State variables for the model. Variables in both columns represent cells or mediators that diffuse between the two compartments.

| Bloodstream | Lung | Description |
| --- | --- | --- |
| | $E_h$ | Healthy epithelial cells |
| | $E_d$ | Damaged epithelial cells |
| | $E_e$ | Dead epithelial cells/empty space |
| $p_b$ | $p$ | Pro-inflammatory mediators |
| $a_b$ | $a$ | Anti-inflammatory mediators |
| $M_{0b}$ | $M_0$ | Unactivated macrophages |
| $M_{1b}$ | $M_1$ | M1 pro-inflammatory macrophages |
| $M_{2b}$ | $M_2$ | M2 anti-inflammatory macrophages |
| $N_{0b}$ | | Unactivated neutrophils |
| $N_b$ | | Activated neutrophils |
| | $N$ | Neutrophils |
| | $AN$ | Apoptotic neutrophils |
| | $R$ | Repair mediators |

**Table 2:** Model parameters with short descriptions and ranges used in LHS.

| Name | Description | Range used |
| --- | --- | --- |
| $a_{b\infty}$ | Relative effectiveness of $a_b$ at inhibiting $M_{0b}$ differentiation to $M_{1b}$ | [0.29, 67.35] |
| $a_{\infty}$ | Relative effectiveness of $a$ at inhibiting $M_0$ differentiation to $M_1$ | [0.13, 72.08] |
| $b_d$ | Baseline decay of damaged cells | $[1.06 \times 10^{-5}, 0.07]$ |
| $b_p$ | Baseline self-resolving repair of epithelial cells | [0, 6.20] |
| $b_r$ | Baseline repair of damaged cells | $[9.79 \times 10^{-3}, 4.47]$ |
| $d_a$ | Rate of diffusion for $a$ | [0.19, 177.98] |
| $d_p$ | Rate of diffusion for $p$ | $[0.34, 2.3 \times 10^3]$ |
| $d_{m0}$ | Rate of diffusion for $M_0$ | [0.24, 275.55] |
| $d_{m1}$ | Rate of diffusion for $M_1$ | $[2.75 \times 10^{-3}, 19.8]$ |
| $d_{m2}$ | Rate of diffusion for $M_2$ | [0.14, 143.36] |
| $k_{am1}$ | Production rate of $a$ by $M_{1b}$ & $M_1$ | [0.01, 18.01] |
| $k_{am2}$ | Production rate of $a$ by $M_{2b}$ & $M_2$ | $[2.43 \times 10^{-3}, 1.67]$ |
| $k_{an}$ | Rate at which neutrophils become apoptotic | [0.01, 50.04] |
| $k_{anm1}$ | Rate of $M_1$ phagocytosis of $AN$ | $[1.32 \times 10^{-3}, 0.69]$ |
| $k_{anm2}$ | Rate of $M_2$ phagocytosis of $AN$ | $[2.71 \times 10^{-3}, 7.36]$ |
| $k_{em1}$ | Rate of phagocytosis of damaged cells by $M_1$ | [0.01, 16.03] |
| $k_{en}$ | Rate of phagocytosis of damaged cells by $N$ | [0.01, 16.03] |
| $k_{ep}$ | Rate of self-resolving repair mediated by $p$ | [0, 4.30] |
| $k_{er}$ | Rate of repair of damaged cells by $R$ | $[1.47 \times 10^{-3}, 1.08]$ |
| $x_{er}$ | Regulates effectiveness of repair of damaged cells by $R$ (Hill-type constant) | $[7.23 \times 10^{-3}, 4.13]$ |
| $k_{m0a}$ | Rate of differentiation of $M_0$ by $a$ | [0.01, 89.07] |
| $x_{m0a}$ | Regulates effectiveness of differentiation of $M_0$ by $a$ (Hill-type constant) | [0.16, 136.83] |
| $k_{m0ab}$ | Rate of differentiation of $M_{0b}$ by $a_b$ | [1.15, 436.59] |
| $x_{m0ab}$ | Regulates effectiveness of $a_b$ differentiation of $M_{0b}$ (Hill-type constant) | [0.16, 83.97] |
| $k_{m0ad}$ | Rate of recruitment of $M_{0b}$ by $a_b$ | [0.34, 181.89] |
| $x_{m0ad}$ | Regulates effectiveness of recruitment of $M_{0b}$ by $a_b$ (Hill-type constant) | [0.01, 27.6] |
| $k_{m0p}$ | Rate of differentiation of $M_0$ by $p$ | $[8.99 \times 10^{-3}, 37.2]$ |
| $x_{m0p}$ | Regulates effectiveness of differentiation of $M_0$ by $p$ (Hill-type constant) | $[1.17, 1.14 \times 10^4]$ |
| $k_{m0pb}$ | Rate of differentiation of $M_{0b}$ by $p_b$ | [0.05, 89.96] |
| $x_{m0pb}$ | Regulates effectiveness of differentiation of $M_{0b}$ by $p_b$ (Hill-type constant) | $[41.51, 2.92 \times 10^4]$ |
| $k_{m0pd}$ | Rate of recruitment of $M_{0b}$ by $p_b$ | $[4.57 \times 10^{-3}, 53.97]$ |
| $x_{m0pd}$ | Regulates effectiveness of recruitment of $M_{0b}$ by $p_b$ (Hill-type constant) | [0.24, 180.74] |
| $k_{m1p}$ | Rate of recruitment of $M_{1b}$ by $p_b$ | [0.2, 92.81] |
| $x_{m1p}$ | Regulates effectiveness of recruitment of $M_{1b}$ by $p_b$ (Hill-type constant) | $[9.8 \times 10^{-3}, 1.69]$ |
| $k_{m2a}$ | Upregulation of $M_{2b}$ recruitment by $a$ | [0.1, 219.93] |
| $x_{m2a}$ | Regulates effectiveness of $M_{2b}$ recruitment by $a$ (Hill-type constant) | [0.08, 94.84] |
| $k_{m2r}$ | Upregulation of $M_{2b}$ recruitment by $R$ | $[3.61 \times 10^{-3}, 20.11]$ |
| $x_{m2r}$ | Regulates effectiveness of $M_{2b}$ recruitment by $R$ (Hill-type constant) | [0.01, 18.70] |
| $k_{man}$ | Rate of $M_1$ switch to $M_2$ by $AN$ | [0.01, 27.08] |
| $k_{mne}$ | Rate of collateral damage to epithelial cells by macrophages and neutrophils | $[1.12 \times 10^{-3}, 5.17]$ |
| $x_{mne}$ | Regulates effectiveness of macrophages and neutrophils to damage epithelial cells (Hill-type constant) | [0.03, 41.06] |
| $k_n$ | Rate of migration of $N_b$ to lung | $[2.39 \times 10^{-3}, 3.54]$ |
| $k_{n0p}$ | Rate of activation of $N_b$ by $p$ | [0.01, 5.58] |
| $x_{n0p}$ | Regulates effectiveness of activation of $N_b$ by $p$ (Hill-type constant) | [0.03, 142.56] |
| $k_{pe}$ | Production rate of $p$ by $E_d$ | $[44.02, 1.12 \times 10^4]$ |
| $k_{pm1}$ | Production rate of $p$ by $M_1$ & $M_{1b}$ | [0.24, 412.22] |
| $k_{pn}$ | Production rate of $p$ and $p_b$ by neutrophils | $[1.67 \times 10^{-3}, 2.95]$ |
| $k_{rm2}$ | Production rate of $R$ by $M_2$ | [0.02, 40.97] |
| $\mu_a$ | Decay rate of $a$ | $[5.16 \times 10^{-4}, 5.08]$ |
| $\mu_{ab}$ | Decay rate of $a_b$ | [0.04, 12.86] |
| $\mu_p$ | Decay rate of $p$ | $[2.76 \times 10^{-3}, 41.04]$ |
| $\mu_{pb}$ | Decay rate of $p_b$ | $[4.79 \times 10^{-4}, 3.71]$ |
| $\mu_{m0}$ | Decay rate of $M_0$ | [0.01, 42.67] |
| $\mu_{m0b}$ | Decay rate of $M_{0b}$ | $[7.66 \times 10^{-3}, 329.59]$ |
| $\mu_{m1}$ | Decay rate of $M_1$ | $[8.2 \times 10^{-3}, 10.16]$ |
| $\mu_{m1b}$ | Decay rate of $M_{1b}$ | [0.03, 60.32] |
| $\mu_{m2}$ | Decay rate of $M_2$ | [0.27, 135.37] |
| $\mu_{m2b}$ | Decay rate of $M_{2b}$ | [0.02, 16.51] |
| $\mu_{nb}$ | Decay rate of $N_b$ | $[2.49 \times 10^{-3}, 6.03]$ |
| $\mu_{n0b}$ | Decay rate of $N_{0b}$ | $[3.94 \times 10^{-6}, 2.1 \times 10^{-3}]$ |
| $\mu_n$ | Decay rate of $N$ | $[8 \times 10^{-3}, 4.32]$ |
| $\mu_R$ | Decay rate of $R$ | [0.72, 761.75] |
| $s_a$ | Source rate of background $a_b$ | $[5.75 \times 10^{-3}, 1.11]$ |
| $s_d$ | Rate of damage from ventilator | 0.75 |
| $s_m$ | Source rate of $M_{0b}$ | $[1.28, 1.14 \times 10^3]$ |
| $s_n$ | Source rate of $N_{0b}$ | [0.22, 225.45] |
| $s_p$ | Source rate of background $p_b$ | $[6.5 \times 10^{-4}, 9.4]$ |

#### 2.3.1. Epithelial cells

We continue with the convention of three subpopulations of epithelial cells, as in Eqs (6) and (7) with *E*_*e*_ = 1 − *E*_*h*_ − *E*_*d*_. We add more details in Eqs (8), (9), and (10) to describe interactions with the immune response variables that we now explicitly model for a more accurate representation of the response to VILI. The first term in Eq (8) is still a logistic growth, representing epithelial cells that spread and replicate to fill *E*_*e*_. This term appears negated in Eq (10), modeling the removal of empty space. The next term in Eq (8) and the first term of Eq (9) represents repair of damaged cells back to a healthy state. Epithelial cells are prone to self-repair [46], represented by a baseline rate *b*_*r*_, and repair at a faster rate in the presence of repair mediators variable *R*, which tracks the level of mediators that promote epithelial repair such as fibronectin and other epithelial growth factors [48, 51, 52]. The third term in Eq (8) and second in Eq (9) represents collateral damage to epithelial cells by the influx and activity of the immune system. This mechanism is modeled via a nonlinear term, which is dependent on macrophage and neutrophil levels [14, 50, 53]. We also model damage due to stretch induced by the ventilator as *s*_*d*_*E*_*h*_, the fourth term in Eq (8) and fifth term in Eq (9), in which injury occurs at a rate proportional to the amount of healthy epithelial cells at a given time.

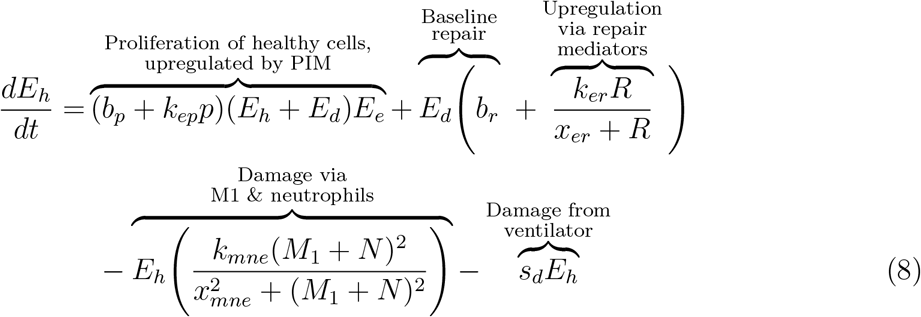

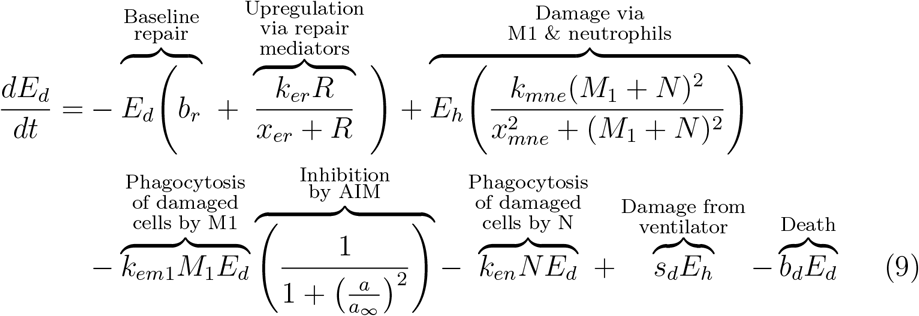

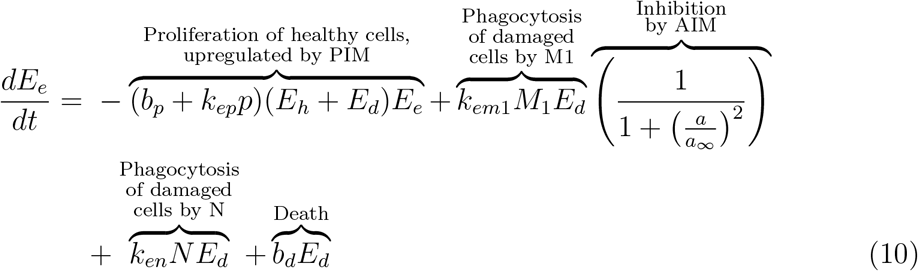

M1 macrophages and neutrophils clear debris from the inflammation site to make room for healthy epithelial cells to divide and fill the empty space [17, 46, 47]. The third and fourth terms in Eq (9) represent this phagocytosis of damaged cells by M1 macrophages and activated neutrophils, respectively. Regulation of M1 is modeled by the last multiplier in the term, representing inhibition by anti-inflammatory mediators (AIM) such as IL-10 [14, 48, 54]. The negative feedback loop of AIM inhibiting further pro-inflammatory functions occurs frequently in our model in a number of equations described below, and we will heretofore refer to this multiplier as inhibition by AIM. Depending on the compartment, the term may utilize the variable *a*_*b*_ (bloodstream) or *a* (local). The anti-inflammatory and regulatory role of M2 macrophages and the balance between M1 and M2 phenotypes is critical for a successful and rapid recovery [16, 48]. The last term of Eqs (9) and (10), *b*_*d*_*E*_*d*_, represents the death of *E*_*d*_ (negative in Eq 9) and the associated gain in the *E*_*e*_ population (positive in Eq 10)).

Dead epithelial cells and “empty” space are grouped together and modeled by the variable *E*_*e*_ in Eq (10). In the epithelial-only model, *E*_*e*_ was modeled as 1 − *E*_*h*_ − *E*_*d*_. Since mass in conserved in these three equations (the sum of terms in the epithelial differential equations is zero), *E*_*e*_ can be modeled either explicitly, as we chose in Eq (10), or in terms of *E*_*h*_ and *E*_*d*_.

#### 2.3.2. Pro- and anti-inflammatory mediators

As a signal to other immune cells, damaged epithelial cells release proinflammatory cytokines and other mediators, including TNF-*α* and matrix metalloproteinases (MMPs) [15, 46, 47]. In our equations, we group these pro-inflammatory mediators (PIM) into two state variables: *p* in the lungs and *p*_*b*_ in the blood. The release of PIM by damaged epithelial cells leads to diffusion of PIM into the bloodstream to recruit additional immune cells [47]. Movement between model compartments is driven by their difference in concentrations in both Eqs (11) and (12). This simple diffusion term will be used for other variables throughout our model.

M1 macrophages produce PIM, which upregulate the activation and migration of macrophages to the site of injury; see the second term in Eqs (11) and (12) [15, 48]. The macrophage population self-regulates by releasing AIM such as IL-10, thus inhibiting further production of PIM [45]. Therefore the term includes the same inhibiting multiplier as in Eq (9). The rate of PIM production by M1 macrophages decreases with increased concentrations of *a*_*b*_.

Neutrophils are also important producers of pro-inflammatory mediators such as TNF-*α*, IL-1, IL-6, LTB4, and chemokines, which stimulate the activation of macrophages toward an M1 phenotype [17, 18, 52, 53, 55]. Low levels of PIM exist in the absence of damage, accounted for by the source term *s*_*p*_, and we also model natural decay of these mediators.

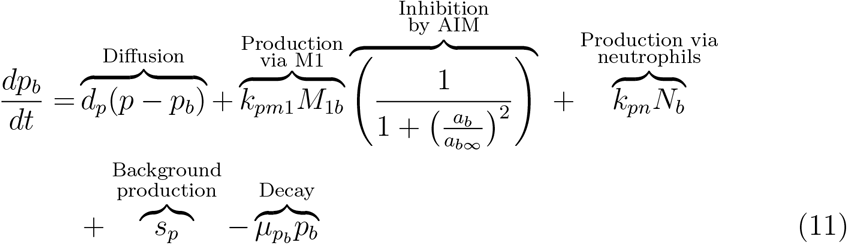

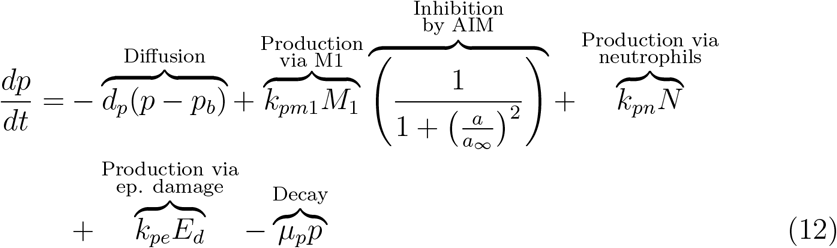

Anti-inflammatory mediators, such as the anti-inflammatory signaling caused by IL-4 and IL-10 [56], are represented by Eq (13) in the blood-stream and Eq (14) at the site of damage. They follow the same simple diffusion behavior as PIM, shown by the first term in each equation below. AIM are released by both M1 and M2 macrophages [15, 48, 54]. Similarly to *p*_*b*_, background levels of *a*_*b*_ are present in the absence of an immune response, represented by term four in Eq (13). Natural decay of AIM is accounted for by the last term in each equation.

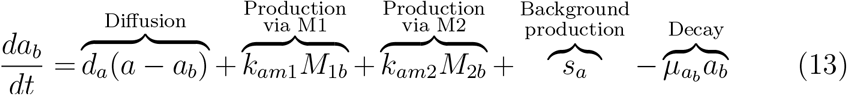

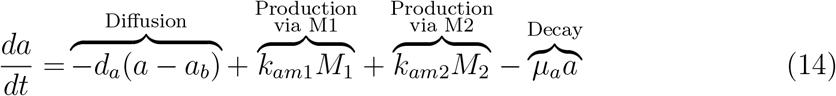

#### 2.3.3. Macrophages

Undifferentiated macrophages, also called naive or unactivated, are present both locally and in the blood. The diffusion term, seen in Eqs (15) and (16), represents movement between compartments. The baseline diffusion between compartments is modeled in the same manner as with other variables, but the rate at which this diffusion occurs is modulated by mediators. Increased PIM and AIM levels cause undifferentiated macrophages in the bloodstream to be recruited at a higher rate to the damaged site, where they become activated and perform phagocytic, pro-inflammatory, and pro-resolving roles [15]. This increased flux between compartments due to the presence of *p*_*b*_ and *a*_*b*_ is modeled by adding to the baseline diffusion rate (*d*_*m*0_). The added term is a Michaelis-Menten-type term to capture the increasing rate as mediators rise, with a maximum rate at which these cells can diffuse, (*d*_*m*0_ + *k*_*m*0*pd*_ + *k*_*m*0*ad*_).

The equations also account for early activation in the bloodstream by PIM and AIM given a high enough concentration of these mediators [14]. Although there is still debate on the types of macrophages that exist in the bloodstream after being released from the bone marrow, there is evidence that populations of both M1 and M2 exist in the bloodstream before being recruited to the site of injury [15, 54]. Thus, we include this process in our equations in the second terms of Eqs (15) and (16). Undifferentiated macrophages in the bloodstream can change phenotype to M1 or M2 after interacting with PIM or AIM, respectively, modeled by a Hill-type term. This nonlinearity accounts for the sufficient amount of PIM or AIM necessary to precipitate activation as well as a saturation of this process.

Once pro-inflammatory mediators such as TNF-*α*, TGF-*β*, and interleukins (ILs) [47] are released by damaged epithelial cells, undifferentiated macrophages receive these signals and differentiate into the M1 phenotype [57]. A pro-inflammatory response characterizes the early stages of the immune response [48, 52]. The second term in Eqs 15 and 16 represent activation of undifferentiated macrophages to the pro-inflammatory phenotype, downregulated by the anti-inflammatory response through an inhibition multiplier. In this term, M2 macrophages can also be activated directly from the naive phenotype by various repair and anti-inflammatory mediators involved in the repair of epithelial cells [47, 48].

Using the same inhibition multiplier as previously, AIM inhibit differentiation to M1 as part of their regulatory role in the inflammatory process, although a complete understanding of these mechanisms is yet to be uncovered [15, 45, 47]. In the absence of injury, lungs contain a low number of undifferentiated macrophages which patrol the surrounding area [46]. “Patrolling” macrophages are also prevalent in the bloodstream. The third term in Eq (15) represents a constant source of undifferentiated macrophages from the circulation [48]. We also account for natural decay of all macrophage phenotypes in Eqs (15) through (20).

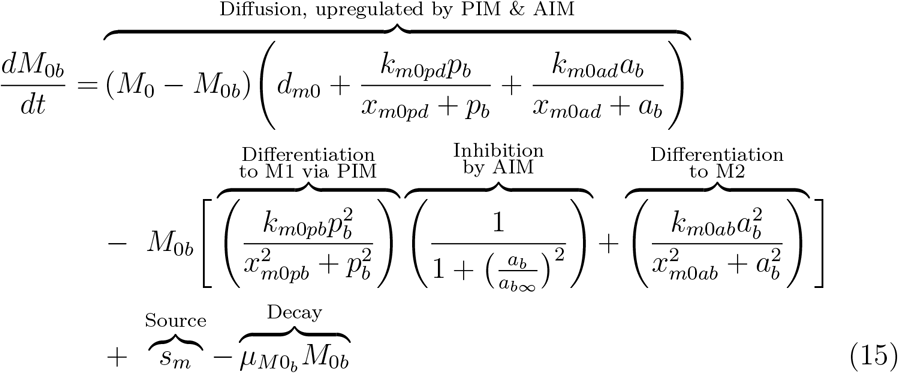

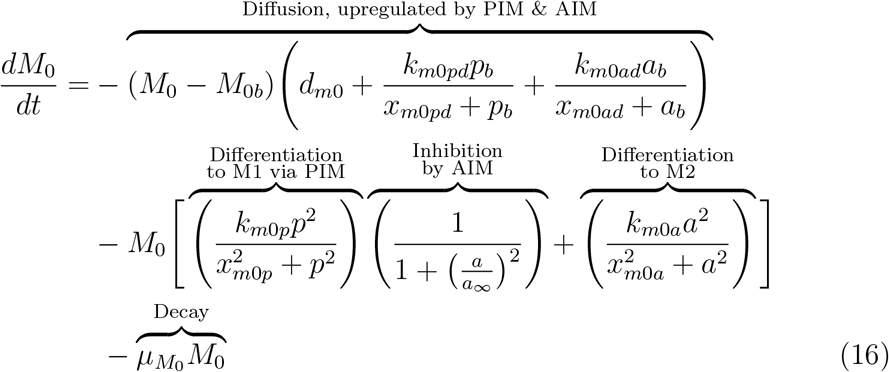

Similarly to naive macrophages, M1 macrophages move between compartments. The presence of pro-inflammatory mediators, which act as recruiters, increases the rate of diffusion, shown in the first term of Eq (17) [15]. The second term represents differentiation from the naive state, as described above.

Macrophages exhibit high plasticity, and based on the mediators and other immune cells they encounter, they can switch phenotype and perform different or enhanced functions; this plasticity is not yet fully understood [14, 48]. M1 macrophages are primarily responsible for producing PIM, thereby recruiting other immune cells to the damaged area [54]. M2 macrophages are considered pro-resolving and downregulate PIM. Both M1 and M2 macrophages phagocytize apoptotic cells such as neutrophils [52]. The shift from an overall pro-inflammatory phase to an anti-inflammatory phase in the course of the immune response is highly dependent upon a shift in macrophage behavior, specifically the shift from a mainly M1 response to a mainly M2 response [15, 47, 54].

One of the primary ways this shift is achieved is through the inhibition of M0 to M1 differentiation by anti-inflammatory mediators, as described previously. Additionally, when pro-inflammatory macrophages phagocytize apoptotic neutrophils, they shift towards a more anti-inflammatory phenotype. This results in suppression of the release of pro-inflammatory mediators and production of pro-resolving mediators [50, 53]. We account for this shift by including the third term in Eq (18), proportional to apoptotic neutrophil phagocytosis which causes M1 macrophages to shift to the M2 phenotype. This term also includes inhibition of M1 function by AIM. It has been shown in some studies that M2 macrophages can switch to an M1 phenotype [58], although this idea is not currently widely accepted. Thus, we choose to include only the shift from M1 to M2.

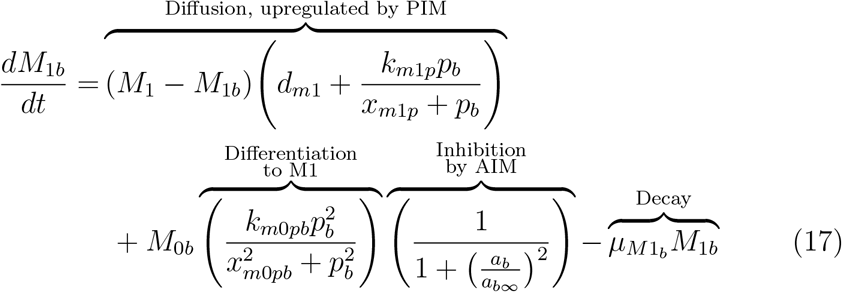

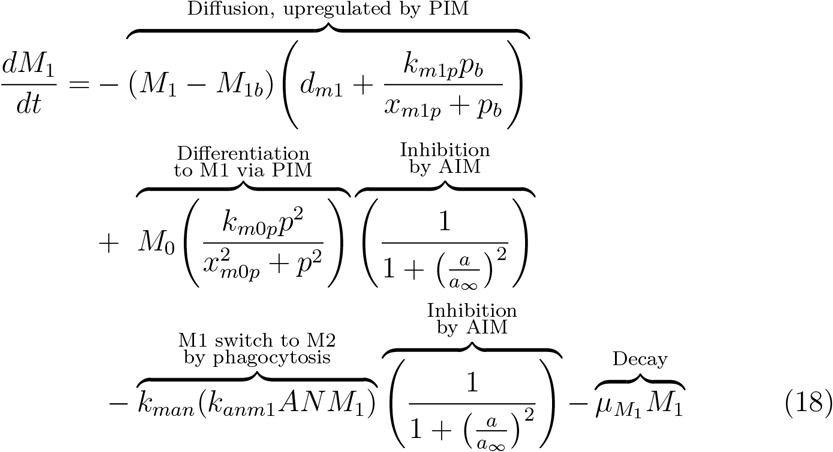

M2 macrophages, associated with an anti-inflammatory response, can be activated directly from undifferentiated macrophages by specific anti-inflammatory signals in addition to switching phenotype from M1. They diffuse between compartments as illustrated previously, shown in the first terms in Eqs (19) and (20). M2 macrophages produce anti-inflammatory mediators which recruit and promote differentiation to more M2 macrophages, described in the second term of both equations. They release cytokines that trigger the repair phase of the immune response [15, 48]. This repair phase includes repair mediators (discussed below in Eq (25)), which play a direct role in the reconstruction of healthy epithelial cells and resolution of damage [48].

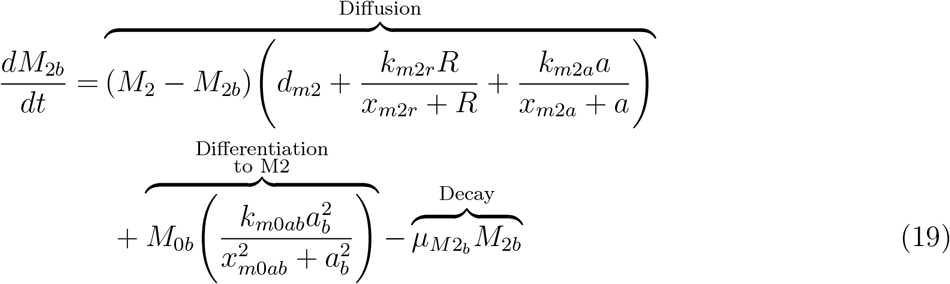

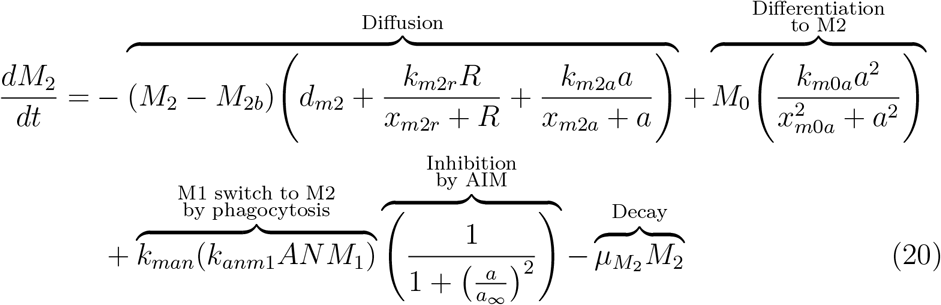

#### 2.3.4. Neutrophils

Neutrophils are considered the first responders to injury [18, 47]. Generated in the bone marrow [17], free-flowing neutrophils circulate in the vasculature at baseline levels, described as *N*_*0b*_ and represented by the first term in Eq (21) [18]. In the presence of injury, neutrophils are activated and recruited to the damaged site through pro-inflammatory mediators such as TNF-*α*, IL-1*β*, and other chemokines and cytokines [18, 55]. This recruitment is represented by the first term in Eqs (21) and (22). On the other hand, anti-inflammatory mediators, including macrophage-produced resolvins and protectins, inhibit further recruitment of neutrophils [50]. Similarly to the differentiation of macrophages, it is assumed that a higher concentration above baseline is required for neutrophils to activate, and that this activation rate saturates. Therefore, a Hill-type term with a maximum rate of *k*_*n0p*_ and a constant of *x*_*n*0*p*_ is used to model activation of neutrophils by PIM. To model the inhibition of neutrophil activation by AIM, we include the same inhibition multiplier as previously described. The effectiveness of these AIMs to inhibit this process is controlled by *a*_*b*∞_. We also account for intrinsic decay of neutrophils in the last term of Eqs (21) through (24).

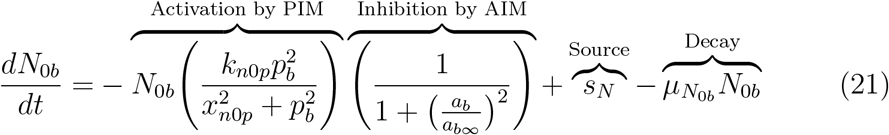

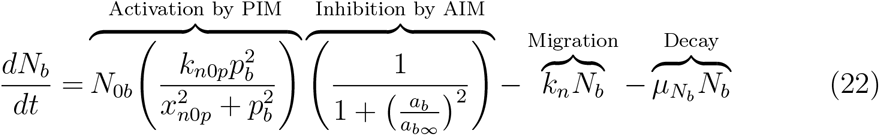

Neutrophils go through a multi-step process of rolling along and subsequently adhering to the surface of the endothelium. Then neutrophils transmigrate to the injury site either through or between endothelial cells [17, 18]. This process is assumed to be driven not by a concentration difference in neutrophils between the compartments but rather is a direct consequence of activation. Therefore, neutrophil transmigration, the first term in Eq (23), is modeled from the bloodstream to the site of injury by a linear term with rate *k*_*n*_.

Activated neutrophils that have transmigrated through the endothelium and reached the site of injury release pro-inflammatory mediators, as discussed previously in Eq (12). During infection, neutrophils play an important role by phagocytizing pathogens [53], but during VILI a main role of neutrophils is the recruitment of macrophages, particularly to promote a more pro-inflammatory environment for the clearance of damaged and dead cells [18].

Neutrophils become apoptotic, modeled by the second term of Eq (23) [47]. In this state, they are phagocytized by M1 and M2 macrophages (second and third terms of Eq (24), respectively) and no longer contribute to the production of PIM [17, 52, 59]. Phagocytosis by M1 macrophages is inhibited by AIM using our standard functional form for the inhibition multiplier. AIM do not inhibit phagocytosis by M2 macrophages since AIM support the function of anti-inflammatory cells. Intrinsic decay is described in the last term of Eq (23).

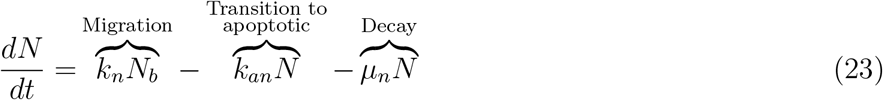

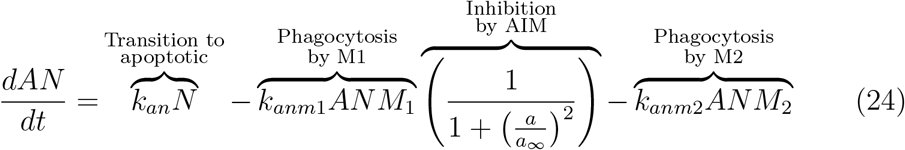

#### 2.3.5. Repair mediators

The direct contribution of alveolar macrophages to the repair of epithelial cells is not completely understood, although macrophage involvement in the repair process has been widely demonstrated [48]. M2 macrophages produce various mediators that promote repair of epithelial cells. We do not model each of these explicitly, instead we group them together in one variable called *R*. These secreted mediators include prostaglandin E_2_, chemokines such as CCL2, TGF-*β*, fibronectin 1 and other epithelial growth factors [48, 51, 52]. The production of *R* by M2 macrophages is modeled by the first term in Eq (25). The second term models intrinsic decay of these mediators.

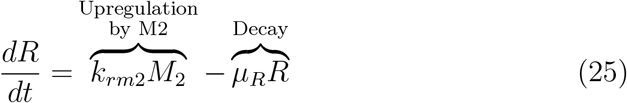

With a system of ODEs that captures the most important aspects of the immune response to VILI, the following sections demonstrate how we analyzed the model to understand the parameter space, determine the most sensitive parameters and other influential predictors of model output, and modulate a particular case of model-generated dynamics to lessen long-term epithelial damage.

### 2.4. Sampling method for parameters: Latin hypercube sampling

Because of the large number of variables and parameters, mathematical and statistical techniques need to be used to analyze the system and find parameter sets that generate biologically realistic dynamics of immune cell populations included in this model. Some parameters may be easily obtained from the literature, such as half-lives of immune cells. However, most of the parameters have not yet been evaluated due to the need for experimental data or are altogether impossible to obtain through current experimental methods. As an initial step towards determining initial conditions and parameters for this model we use Latin hypercube sampling (LHS). Introduced in 1979 [60], LHS is a sampling method which generates random, unique parameter sets, such that the produced parameter values are selected according to a distribution; in our case, a uniform distribution. For LHS with uniform distributions assumed for each parameter, to generate *n* desired parameter sets, the algorithm splits the determined range into *n* evenly-spaced subintervals and each interval is sampled exactly once [61]. This is particularly useful for our exploratory simulations because the distributions of the parameters are unknown.

Using MATLAB functions adapted from Kirschner *et al.* [62], all parameters were sampled except the rate of damage *s*_*d*_ due to ventilation. We used simulations to explore parameter space by sampling near transients associated with different types of disease progression. We accepted parameter sets if they were associated with a steady state solution and defined the final parameters ranges based on the maximum and minimum value of the parameters in the acceptable sets. See Table 2 for ranges used for each parameter. Using LHS with these ranges we generated 100,000 parameter sets. Future work could calibrate cohorts to data from different experimental or clinical groups and then use the analysis methods here to compare dynamics and parameters that drive differences between experimental or clinical groups.

### 2.5. Cohorts: Healthy, Moderate Inflammation, & Severe Inflammation

We needed to start our simulation from initial conditions associated with a steady state, so that when ventilation was simulated we were seeing changes in the dynamics only due to the ventilator. For all 100,000 parameter sets we ran the model for 800 hours without ventilator-induced damage (*s*_*d*_ = 0) using three different initial conditions to determine if a steady-state condition was reached in the absence of ventilation. The first initial condition was related to the initial simulations used to develop the sampling ranges and gave rise to 25,195 sets that reached steady state. Additionally, we checked whether parameter sets that did not reach a steady state from these initial conditions could reach a steady state from an initial condition with all variables set to zero except for *E*_*h*_(0) = 0.75 and *E*_*d*_(0) = 0.25 (starting with damaged tissue and no immune response) or initial conditions with all variables set to zero except for *M* 1(0) = 50 (starting with an activated immune response and healthy tissue). These other initial conditions added another 1,104 sets that reached a steady state, bringing the total to 26,299. Any parameter sets that did not result in an equilibrium state by 800 hours from these three initial conditions were not simulated with ventilation. We simulated these 26,299 parameter sets with ventilator-induced damage starting from their steady state levels. Simulations were run for 200 hours with ventilation for the first two hours (a nonzero damage rate), a duration comparable with murine experiments [63, 64].

Many of these sets had initial conditions associated with a severely inflamed lung without ventilation, which did not seem biologically realistic. To correct for this we eliminated sets based on their initial condition for *E*_*e*_ (empty/dead cells). We performed all of the analysis below with three different thresholds to see whether the exclusion of these parameter sets affected the results. In this paper we focus on the 23,086 parameters sets that had *E*_*e*_(0) < 50% and show a summary of all results for *E*_*e*_(0) < 25% and *E*_*e*_(0) < 75% in the supplementary materials. We did not find any major differences when varying this inclusion threshold.

Simulations were separated into three categories of disease progression: 1) healthy epithelial cells sufficiently cover the alveoli to functional normally or existence of 2) moderate or 3) severe inflammation and associated tissue damage. These progressions are called healthy, moderate inflammation, and severe inflammation, respectively.

To quantify these three different states, we divided percentages of healthy epithelial cells into categories:

- Healthy: *E*_*h*_ ≥ 90%
- Moderate inflammation: 50% ≤ *E*_*h*_ < 90%
- Severe inflammation: 0% ≤ *E*_*h*_ < 50%

In this way, each parameter set can be classified into three different categories based on their *E*_*h*_ values either before or after ventilation. Thus, sets are classified by their initial conditions and then again after simulation with ventilation. These parameter sets, their corresponding transients, and the outcomes they generate were used to develop a virtual cohort representing a variety of immune system dynamics. The cohort was then used to compare outcomes, transient properties, underlying parameters, and their corresponding biological mechanisms.

### 2.6. eFAST

We used several tools to perform a sensitivity analysis of model parameters. A common method is calculating partial rank correlation coefficients (PRCCs), but results are only reliable for monotonic relationships between parameters and variables. Our model output does not fit this criteria. Marino *et al.* suggest the extended Fourier amplitude sensitivity test (eFAST), a variance-based method for non-linear, non-monotonic relationships [61]. The greatest drawback of eFAST compared to PRCC is the computation time.

eFAST, developed by Saltelli *et al.* [65], Saltelli & Bolado [66], and Saltelli *et al.* [67] is the extended version of FAST, originally developed by Cukier *et al.* [68], Schaibly & Shuler [69], and Collins & Avissar [70]. Parameters are varied and the resulting variation in model output is calculated using statistical variance. The algorithm varies each parameter at different frequencies by creating a sinusoidal function, called a search curve, and then sampling parameter values along the function. Fourier analysis measures the influence of the parameter’s frequency on model output. First-order sensitivity *S*_*i*_ for a parameter *i* is calculated by varying only *i* and leaving the rest constant. Total-order sensitivity *S*_*Ti*_ is calculated by varying *i* using a unique, higher frequency and varying the other parameters using lower non-unique frequencies. This total-order sensitivity captures non-linear interactions between parameters in addition to changes in model output. We implement the method by Marino *et al.* [61] to calculate *S*_*i*_ and *S*_*Ti*_ and determine their statistical significance of for each parameter. A “dummy parameter” is included in the parameter set and its eFAST index is compared to the other parameters found in the model.

MATLAB functions by Kirschner *et al.* [62] are available online to perform eFAST. We obtain 65 values of each parameter on a search curve and repeat this process for five unique search curves since different ones can generate slightly different samples. Sensitivity can be calculated at specific time points for the desired variable.

### 2.7. Random forest decision tree

Aside from more conventional sensitivity analysis measures, we chose a few alternative methods that require less computation time and can include other features of the model besides parameters. One of these alternatives is a random forest decision tree. A decision tree algorithm is a classification tool that uses the given properties of an individual or object to determine into which category it should fall [71, 72]. In this case, each parameter set in the virtual cohort has a number of predictors and outputs: parameters and any other characteristics from the transients that can be quantified or given a classification value. The algorithm takes a training set, a subset of the cohort about which all predictors and outputs are known, and can train the algorithm to classify virtual cohort members into specific categories.

An output of the model that we are particularly concerned with predicting is the patient’s outcome, as described in the previous section. The decision tree generated from the training set makes predictions for the rest of the virtual cohort members about whether each one will fall into one of the three outcomes: healthy, moderate inflammation, or severe inflammation. The tree contains branches at which specific parameters are chosen to best assist in classification. The parameter values of each “individual” in the cohort determines the path along the tree until it reaches the most likely outcome based on the training set.

Since a decision tree simply takes a series of values for each predictor and is not dependent on the model itself, measures besides just parameters can be used. We included supplementary predictors calculated from the transients, described in Table 3. Adding these predictors allowed for the possibility that the best classifiers of outcome could be not only parameters but also properties of the transients. This knowledge could provide additional information about metrics for experimentalists and clinicians to keep track of and identify early warning signs for undesirable results.

**Table 3:** Additional predictors used in analysis of parameter space with descriptions if necessary. These predictors were used with the random forest decision tree, correlations, and significance testing.

| Predictor | Comment, description |
| --- | --- |
| Maximum $M1$ percent | |
| Maximum $M2$ percent | |
| Minimum $M1$ percent | |
| Minimum $M2$ percent | |
| Maximum $M1$ | |
| Maximum $M2$ | |
| Minimum $M1$ | |
| Minimum $M2$ | |
| $M1$ peak time | Time at which $M1$ peak occurs |
| $M2$ peak time | Time at which $M2$ peak occurs |
| $M2$ percent at 10 hours | |
| $M1$ peak ratio | Ratio of $M1$ peak to $M1$ initial condition |
| $E_h$ difference | Difference between first and last time points of $E_h$ |
| $E_h$ ratio 0.5h | Ratio of IC to $E_h$ at 30 minutes |
| $E_h$ ratio 2h | Ratio of IC to $E_h$ at 2 hours |
| $E_h$ ratio 6h | Ratio of IC to $E_h$ at 6 hours |
| Fits $t = 0$ $M0$ data | 0 = does not fit, 1 = does fit |
| Fits all data | 0 = does not fit, 1 = does fit |

For added robustness against overfitting [72], we use a random forest decision tree algorithm, in which a user-specified number of randomly chosen parameters are candidates at each branch; then the algorithm selects one to be the splitting variable from that smaller group. The `rf` function in R generates 500 decision trees as the “forest” along with several other useful output metrics. One metric in particular is the importance value of each parameter or characteristic, calculated from the Gini Index. The importance value is a measure of how important any given parameter was in determining the outcome of each parameter set in the virtual cohort. Because of the large number of parameters in the model, this can provide intuition on which parameters and other characteristics of the transients are most influential in determining outcomes. The R and MATLAB code used for this method are provided in the supplementary materials.

## 3. Results

Our aim is to understand how recruitment of the immune response and its interactions with epithelial cells translate to specific outcomes and what dynamics are driving this process. Therefore, we developed an ODE model of the immune response to ventilator-induced damage, which explicitly tracks macrophage phenotype and epithelial cells. A fixed point and stability analysis of the epithelial subsystem reveals the long-term stability of a simplified version of the system under various conditions, and how changes in those conditions affect stability. Using Latin hypercube sampling, we generated parameter sets that replicate different possible responses to VILI and created a virtual cohort of patients. We also perform an analysis of the large parameter space by comparing various techniques to determine predictors of outcome and/or processes that could be targeted to modulate outcome.

### 3.1. Sample Transients and Cohort Breakdown

This model can generate a variety of dynamics, similar to expected responses of patients on a ventilator. There is significant variability between outcomes as well as within them. Fig 4 shows examples of these different dynamics for healthy epithelial cells and M0, M1, and M2 macrophages using a case of each of the three outcomes: healthy, moderate inflammation, and severe inflammation. Simulations were run in MATLAB using the code provided in the supplementary materials.

**Figure 4:**
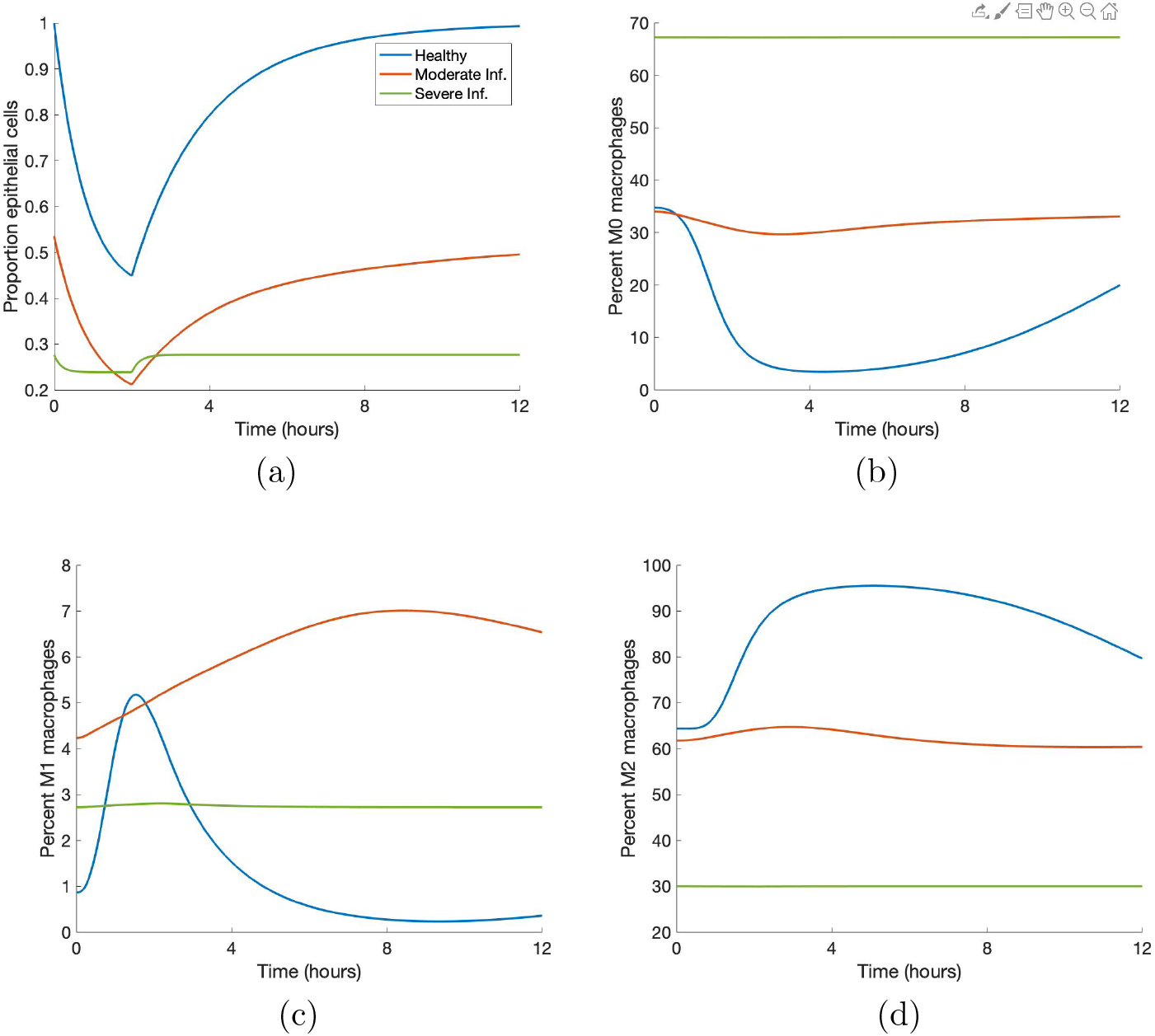
Sample simulations show the variety of model-generated dynamics. Blue, orange, and green curves indicate healthy, moderate inflammation and severe inflammation outcomes, respectively. (a) Proportion healthy epithelial cells. (b) Percent M0 macrophages. (c) Percent M1 macrophages. (d) Percent M2 macrophages.

We generated 100,000 parameter sets using LHS with parameter ranges given in Table 2. Fig 5 shows the breakdown of these parameter sets based on whether or not the dynamics lead to a steady-state system in the absence of ventilation, their classification before ventilation, and the resulting state (healthy, moderate inflammation, and severe inflammation) after 200 hours, the first 2 hours being ventilation. We also rejected any parameter sets with *E*_*e*_(0) ≥ 50%, since this would not be biologically realistic. The top number in each box is the total number of parameter sets in that category, and that number is further broken down by the category in which they start (column 1) and end (column 2). For the first column, the number in parentheses is the number of sets that started in that category but ended in a different one. Conversely, the number in parentheses in the second column shows the sets that ended in a certain outcome but did not start there. These numbers serve as a summary of how damage may affect outcome for the variety of behaviors in the virtual cohort. We will analyze all 23,086 sets that reach steady state (with *E*_*e*_(0) < 50%) to understand the full array of responses that could occur. In the future, experimental data could help narrow down responses.

**Figure 5:**
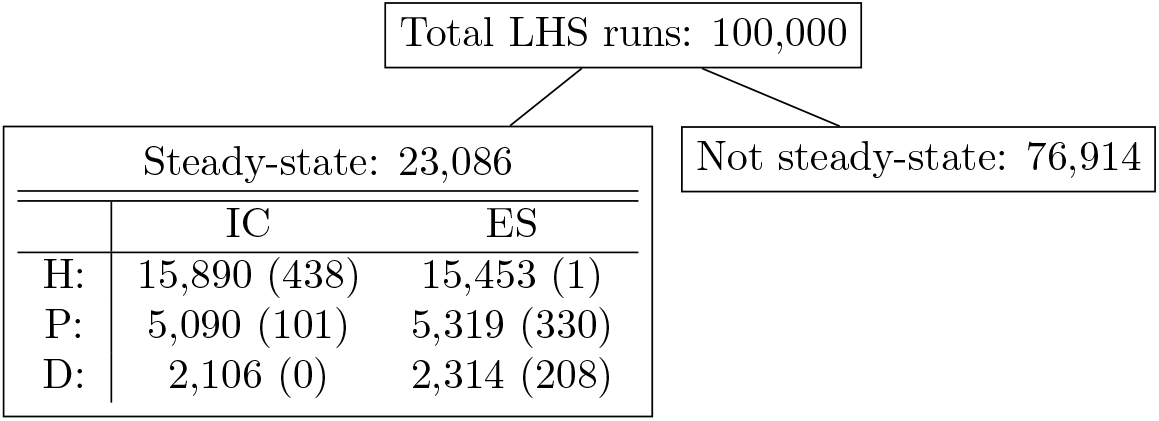
Results of 100,000 LHS runs grouped by disease progression. Parameter sets are broken down by their initial conditions (IC) and ending states (ES) and by category healthy (H), moderate inflammation (M), or severe inflammation (S). Numbers in parentheses in the IC columns are the number of simulations that started in the category associated with that row and change their state after ventilation. Numbers in parentheses in the ES columns are the number of simulations that ended in the category associated with that row, but were not in that category before ventilation. All parameter sets are associated with a steady-state solution with *E*_*e*_(0) < 50%.

### 3.2. Determining Predictors and Driving Dynamics

Our model has 18 variables and 67 parameters. Using a variety of mathematical, statistical, and computational methods, we determined the parameters and other predictors that stand out, those to which output is most sensitive and may help differentiate or predict what is driving outcome. In this section we explain and compare the results of each method.

#### 3.2.1. Correlations and significance testing highlight specific parameters

As an initial step towards understanding relationships between parameters and model output, we calculated the correlations of parameters and predictors with outcome. There were some correlations between predictors that are very high, but are measuring similar things; for example, maximum M1 and minimum M1. We excluded these since they do not provide new or useful information. Aside from these, there are only a few correlations between parameters or between parameters and predictors that are higher than *R* = 0.3; notable pairs are shown in Fig 6 using random samples from each outcome for better visibility of the points. For *k*_*mne*_, the rate of collateral damage to epithelial cells by macrophages and neutrophils, parameter sets that result in moderate and severe inflammation outcomes have a significant correlation with the *E*_*h*_ ratio at 0.5 hours, shown in Fig 6a. The *E*_*h*_ ratio and *k*_*mne*_ have the following correlations for each outcome: healthy *R* = 0.1 (not shown), moderate inflammation *R* = 0.67, and severe inflammation *R* = 0.82. The *b*_*r*_ parameter, representing the baseline repair rate for epithelial cells, has the following correlations with the same *E*_*h*_ ratio for each type of outcome, healthy *R* = 0.29, moderate inflammation *R* = 0.41, and severe inflammation *R* = 0.37, shown in Fig 6b. Visual inspection of both graphs shows possible nonlinear behavior that should be investigated further. The only other pair with a correlation above 0.3 is *s*_*m*_, the source rate for naive macrophages, and the maximum and minimum values of M2 macrophages over the entire simulation. The parameter *s*_*m*_ and maximum M2 have the following correlations: healthy *R* = 0.32; moderate inflammation *R* = 0.3; severe inflammation *R* = 0.3. Fig 6c shows these correlations; *s*_*m*_ and minimum M2 is not shown but have similar results.

**Figure 6:**
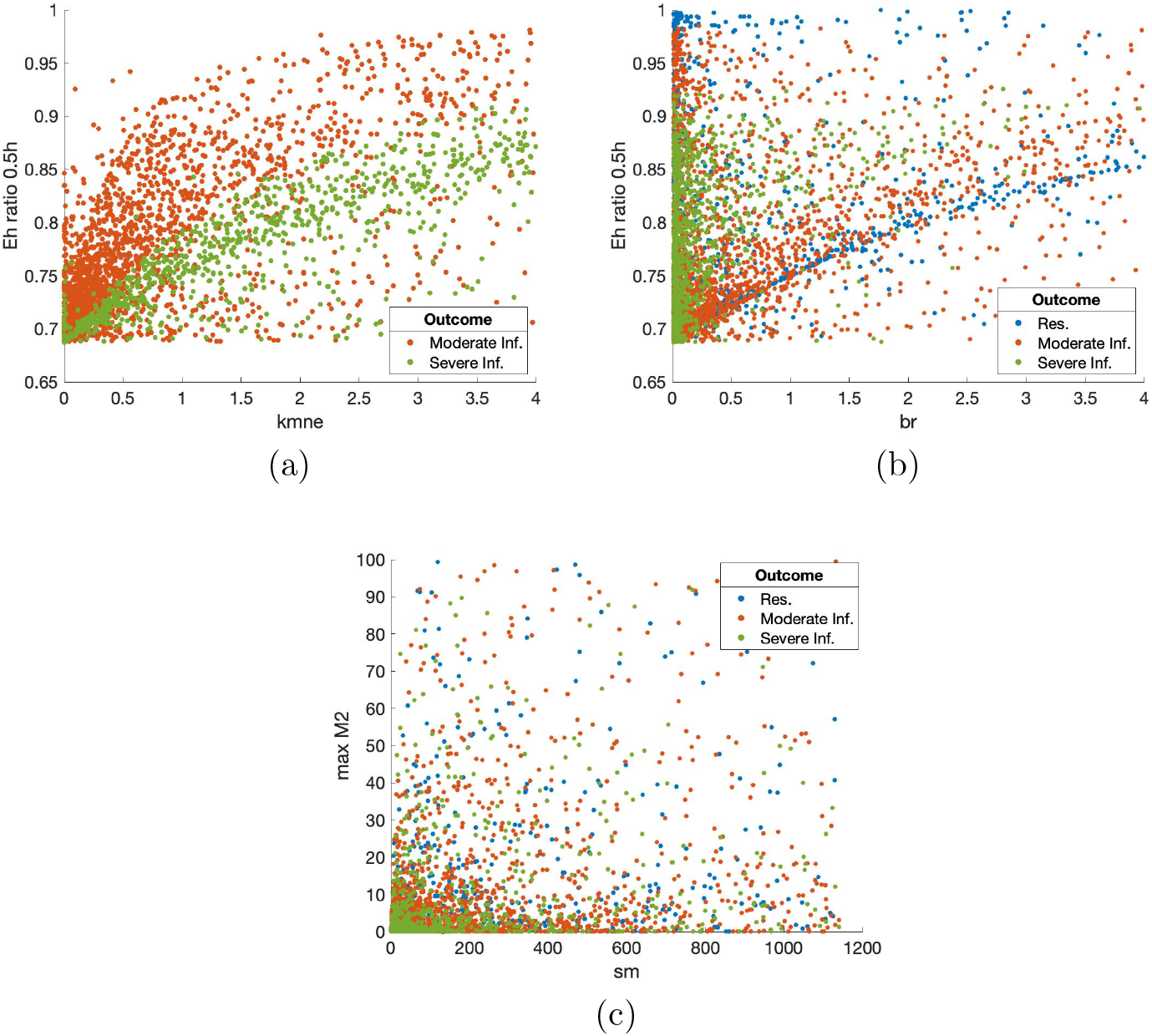
Scatter plot of predictors with notable correlations. Points are a random sample of the total points. (a) Parameter *k*_*mne*_ (rate of collateral damage to epithelial cells by macrophages and neutrophils) versus ratio of *E*_*h*_ at 0.5 hours to initial *E*_*h*_ values. Correlations: resolved to healthy *R* = 0.1 (not shown); moderate inflammation *R* = 0.67; severe inflammation *R* = 0.82. (b) Parameter *b*_*r*_ (baseline rate of epithelial repair) versus ratio of *E*_*h*_ at 0.5 hours to initial *E*_*h*_ values. Correlations for parameter sets in each outcome: resolved to healthy *R* = 0.29; moderate inflammation *R* = 0.41; severe inflammation *R* = 0.37. (c) Parameter *s*_*m*_ (source rate of M0 macrophages) versus maximum M2. Correlations for parameter sets in each outcome: healthy *R* = 0.32; moderate inflammation *R* = 0.3; severe inflammation *R* = 0.3.

We also performed hypothesis testing for predictors (excluding binary variables). The Kruskal-Wallis test is an alternative to ANOVA when the variable distributions are not normal [73]. Due to our choice of a uniform sampling distribution for LHS, parameter distributions for the 23,086 sets are roughly uniform. We categorized all parameter sets by their outcome (healthy, moderate inflammation, severe inflammation) and compared them. If any of the three groups had a statistically significant difference (p-value less than 0.01), a Wilcoxon test was performed on each pair (healthy and moderate inflammation, healthy and severe inflammation, moderate and severe inflammation) to determine which groups were different from one another. P-values for the Kruskal-Wallis and Wilcoxon tests were adjusted using the Benjamini–Hochberg procedure to control for the false discovery rate [74]. Knowledge of which parameters and other predictors are different between groups based on outcome provides insight into predicting outcomes and which predictors might best influence the immune response to damage.

35 out of 81 parameters and predictors returned results for a statistically significant difference between at least two groups and 14 gave statistically significant differences between all three groups. Table 4 shows a summary of the results from the various methods used to examine predictors’ significance in determining model output. Column 1 of Table 4 shows the predictors in which all three groups were different from one another, as determined by the Kruskal-Wallis and Wilcoxon tests. Results in columns 2-5 are described in the following sections. Box plots of a subset of predictors in which all three groups are different are shown in Fig 7 to help visualize these differences.

**Table 4:** Summary of three different methods used to determine the most influential predictors, including parameters and other factors. Columns 1 & 2 show results for all 23,086 parameter sets. Column 1: significance testing results for predictors in which all three outcome groups are statistically different (p-value < 0.01). For ease of comparison between columns, the predictor is listed next to its counterpart in the ordered random forest list, if listed in that column. Column 2: average importance values determined by random forest decision trees. The top ten are ordered from highest to lowest importance. Columns 3-5: eFAST results (ordered by p-value, with p-value < 0.02) for three time points.

| Sig. Testing<br>(Not ordered) | Random Forest<br>(Ordered output) | eFAST (Ordered) |  |  |
| --- | --- | --- | --- | --- |
|  |  | 0.5h | 2h | 6h |
| $k_{mne}$ | $k_{mne}$ | $k_n$ | $k_n$ | $x_{m0a}$ |
| $x_{mne}$ | $x_{mne}$ | $\mu_p$ | $x_{mne}$ | |
| $E_h$ ratio 6h | $E_h$ ratio 6h | $x_{m0a}$ | $k_{en}$ | |
| | $E_h$ ratio 2h | $x_{mne}$ | $b_r$ | |
| | $E_h$ ratio 0.5h | $k_{en}$ | $x_{nup}$ | |
| $b_r$ | $b_r$ | $b_r$ | $x_{m0a}$ | |
| Min M1 | Min M1 | $\mu_{m1}$ | $s_p$ | |
| $k_{en}$ | $k_{en}$ | $k_{am1}$ | $\mu_p$ | |
| Min M1% | Min M1% | | $k_{pe}$ | |
| | $k_{ep}$ | | $\mu_R$ | |
| M1 peak time |  |  |  |  |
| $k_{em1}$ | | | | |
| M2 peak time |  |  |  |  |
| $k_{an}$ | | | | |
| $k_{ep}$ | | | | |
| M1 peak ratio |  |  |  |  |
| $x_{nup}$ | | | | |

**Figure 7:**
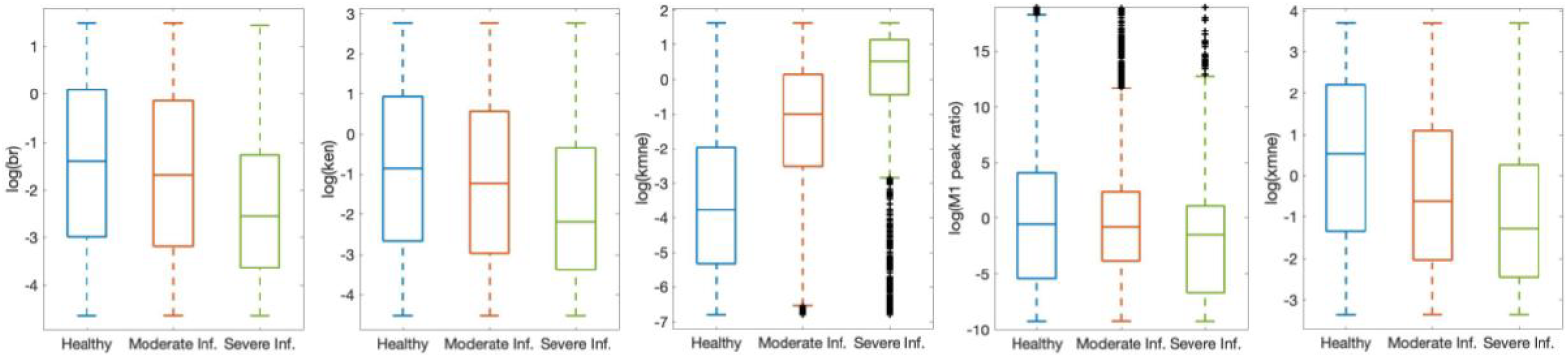
Predictors selected by significance testing show visible differences between disease progression groups. Subset of parameters and predictors that showed a statistically significant difference between all three outcomes: healthy, moderate inflammation, and severe inflammation, as determined by the Kruskal-Wallis and Wilcoxon tests. Some are shown on a log scale and some outliers removed from figure for better visibility. Black x’s are outliers.

#### 3.2.2. Parameter Sensitivity with eFAST

Since outcome of *E*_*h*_ is the metric by which we determine health of the individual, we calculated eFAST indexes for *E*_*h*_ at 30 minutes, two hours (end of ventilation), and six hours. We calculated first-order and total-order sensitivities *S*_*i*_ and *S*_*Ti*_, respectively. Fig 8 shows results for the parameters with p-value < 0.02. Parameters *k*_*n*_ (rate of migration of *N*_*b*_ to lung), *x*_*mne*_ (Hill-type constant for effectiveness of macrophages and neutrophils in damaging epithelial cells), *x*_*m*0*a*_ (Hill-type constant for effectiveness of differentiation of *M*_0_ by *a*), *b*_*r*_ (baseline repair of damaged cells), and *k*_*en*_ (phagocytosis of damaged cells by *N*) are sensitive for several time points. Comparing *S*_*i*_ and *S*_*Ti*_ in Fig 8, it is possible that nonlinear interaction between parameters affects model output more at 6 hours than at 2 hours. Parameters with a significant *S*_*i*_ may also be better candidates for treatment than those with a significant *S*_*Ti*_ because first-order sensitivity measures sensitivity of *E*_*h*_ based only on fluctuations in a single parameter. For this reason and since many of the same parameters are significant for first-order and total-order sensitivity, we show results for first-order sensitivity in Columns 5-7 of Table 4, ordered from lowest p-value to highest and for the three time points specified.

**Figure 8:**
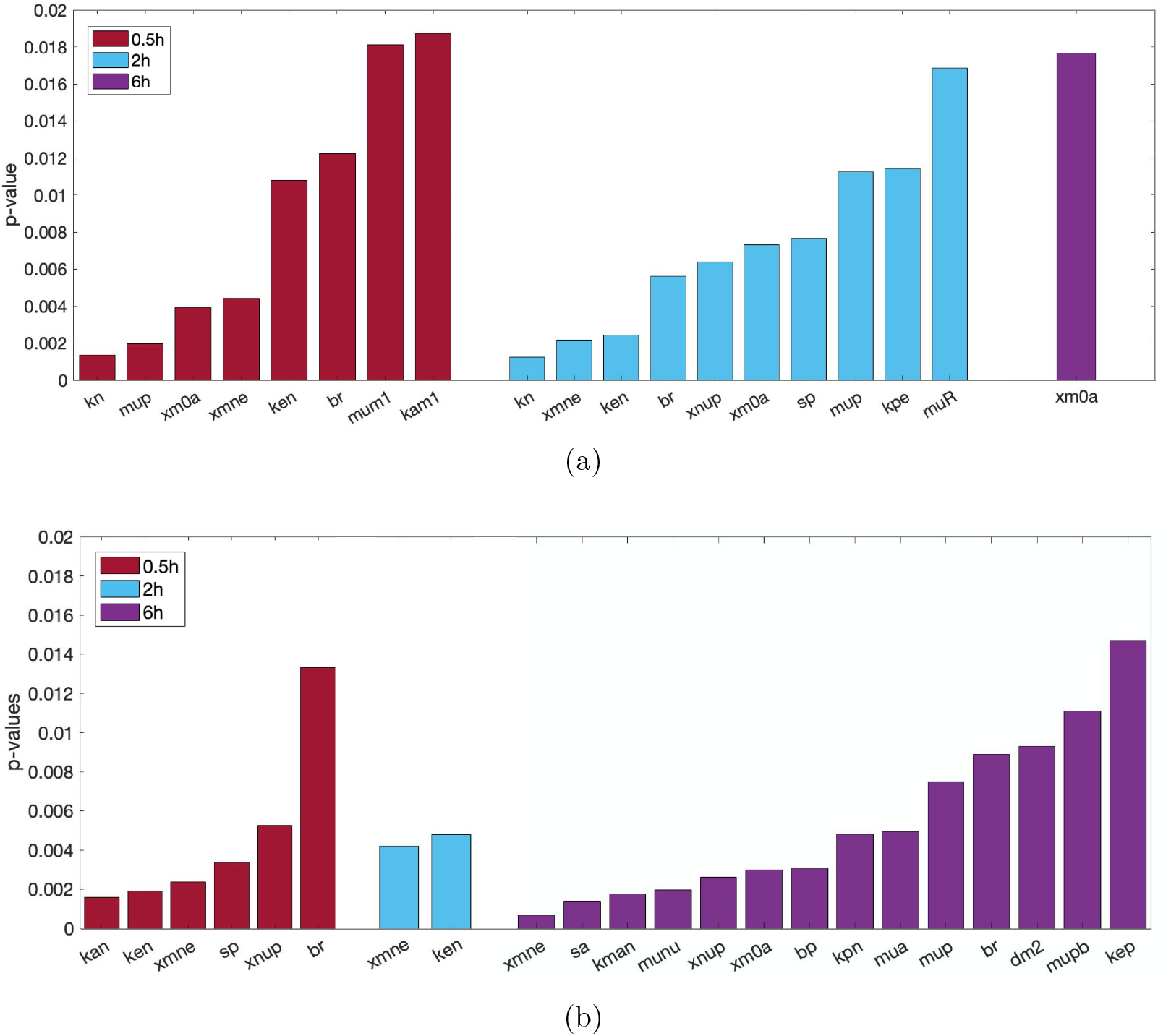
Parameter sensitivity analysis shows which parameters most influence model output. Parameters determined by eFAST to be most sensitive, with p-values calculated by comparing eFAST sensitivity indexes to a dummy variable. Results are given for each of the time points tested: 0.5 (red), 2 (blue), and 6 hours (purple). (a) First-order sensitivity, also shown in Table 4. (b) Total-order sensitivity.

#### 3.2.3. Random forest algorithm to determine predictors

The randomness of the decision tree algorithm means that each random forest generated and its resulting importance values are slightly different. To offset any unusual results generated by the randomness, we replicated the process of randomly selecting a training set and generating importance values from the random forest 1000 times. Fig 9 shows the average and standard deviations of the top ten importance values generated.

**Figure 9:**
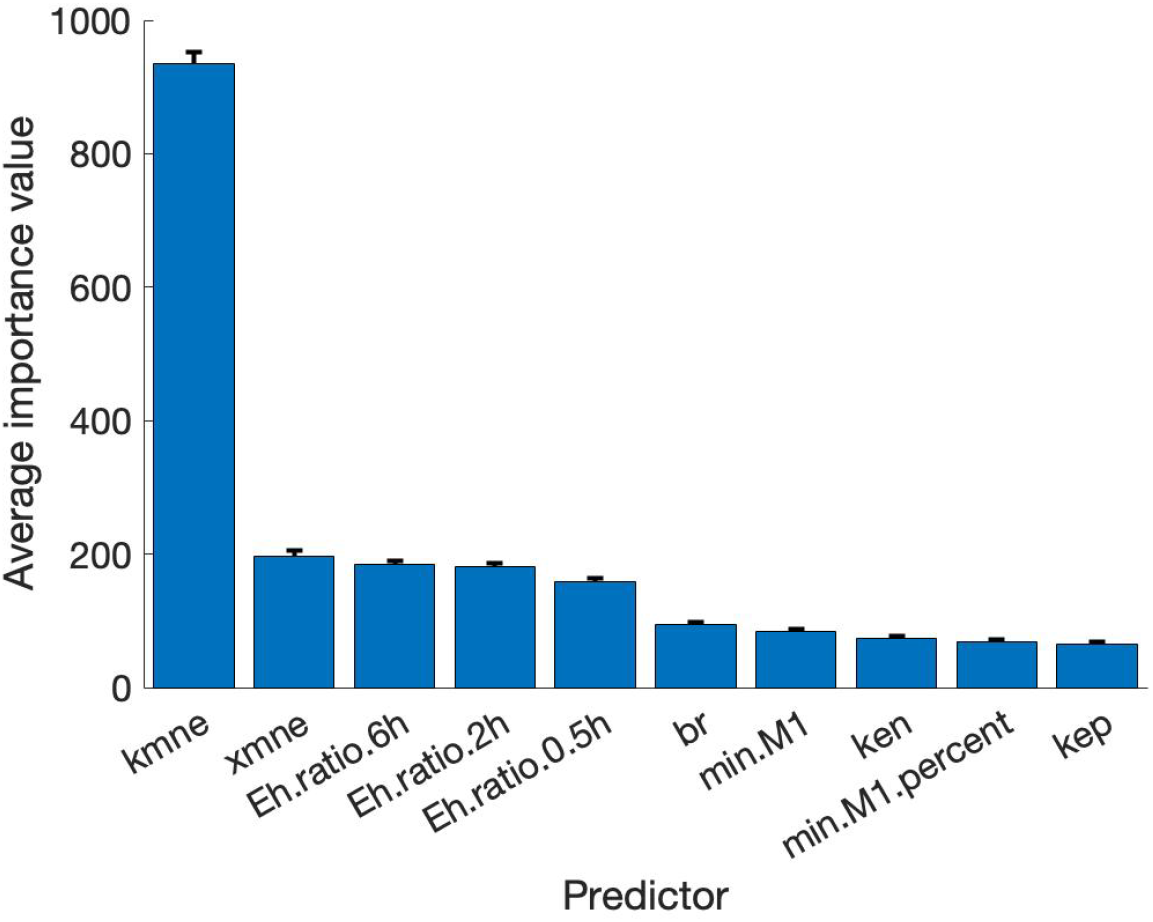
Random forest decision tree selects top indicators of outcome. Mean and standard deviation of importance values for the top ten highest predictors from 1000 random forest decision trees.

Notice that the standard deviations are small enough so that although some of the top importance values may change order in different random forest simulations, in general the most important predictors remained the same across numerous simulations. Furthermore, several of the top ten predictors were found to be significant by the Kruskal-Wallis Test, and *b*_*r*_ and *k*_*mne*_ are shared by random forest and eFAST. (see Table 4). The consistency of the importance of these parameters and predictors using different methods supports the idea that they play a significant role in the sensitivity of model output and determining or differentiating outcomes.

### 3.3. Modulating recovery: a case study of select transients

Fig 10 shows nine examples of transients that started in one disease progression category and ended in another. We used the information gained in the parameter analysis to identify key targets for treatment that could modulate damage, especially in the case of a patient starting in one state and ending in a different, negative outcome after ventilation. The goal is to return the cohort member to its original steady-state earlier, since the inability to recover from a 2-hour vent after 200 hours or more could be detrimental to long-term health.

**Figure 10:**
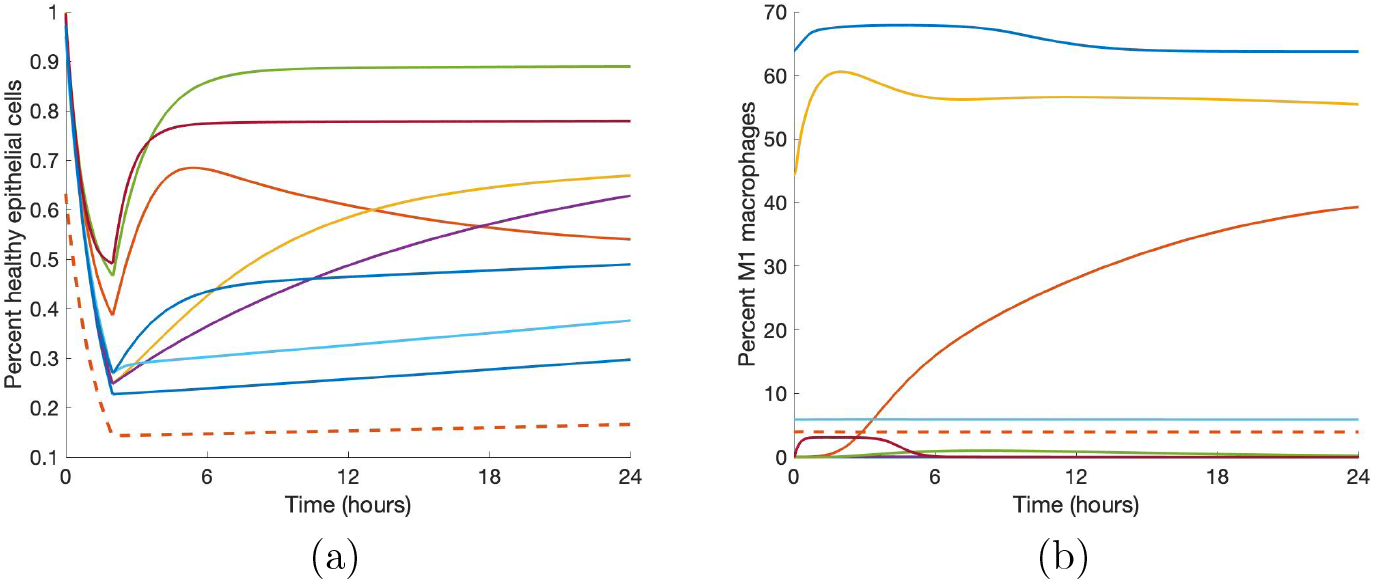
Some parameter sets generate transients that end in a worse disease progression after ventilation. (a) Transients of *E*_*h*_ that start at one state and end at a lower one. (b) Corresponding transients of *M* 1. Solid lines represent transients that start healthy and end in moderate inflammation; the dotted line represents the transient that starts in moderate inflammation and ends in severe inflammation.

Our analysis shows that the parameters *b*_*r*_, the rate of self-repair of healthy epithelial cells, *k*_*mne*_, the rate of collateral damage by macrophages and neutrophils to epithelial cells, *x*_*mne*_, the Hill-type constant which regulates the effectiveness of macrophages and neutrophils in damaging epithelial cells, and *k*_*en*_, the rate of phagocytosis of damaged cells by neutrophils, are some of the most influential parameters and thus could inform targets for treatment. It is also important to note that different interventions could begin and end at any time during or after ventilation, so we examined interventions at several time points (see Fig 11).

**Figure 11:**
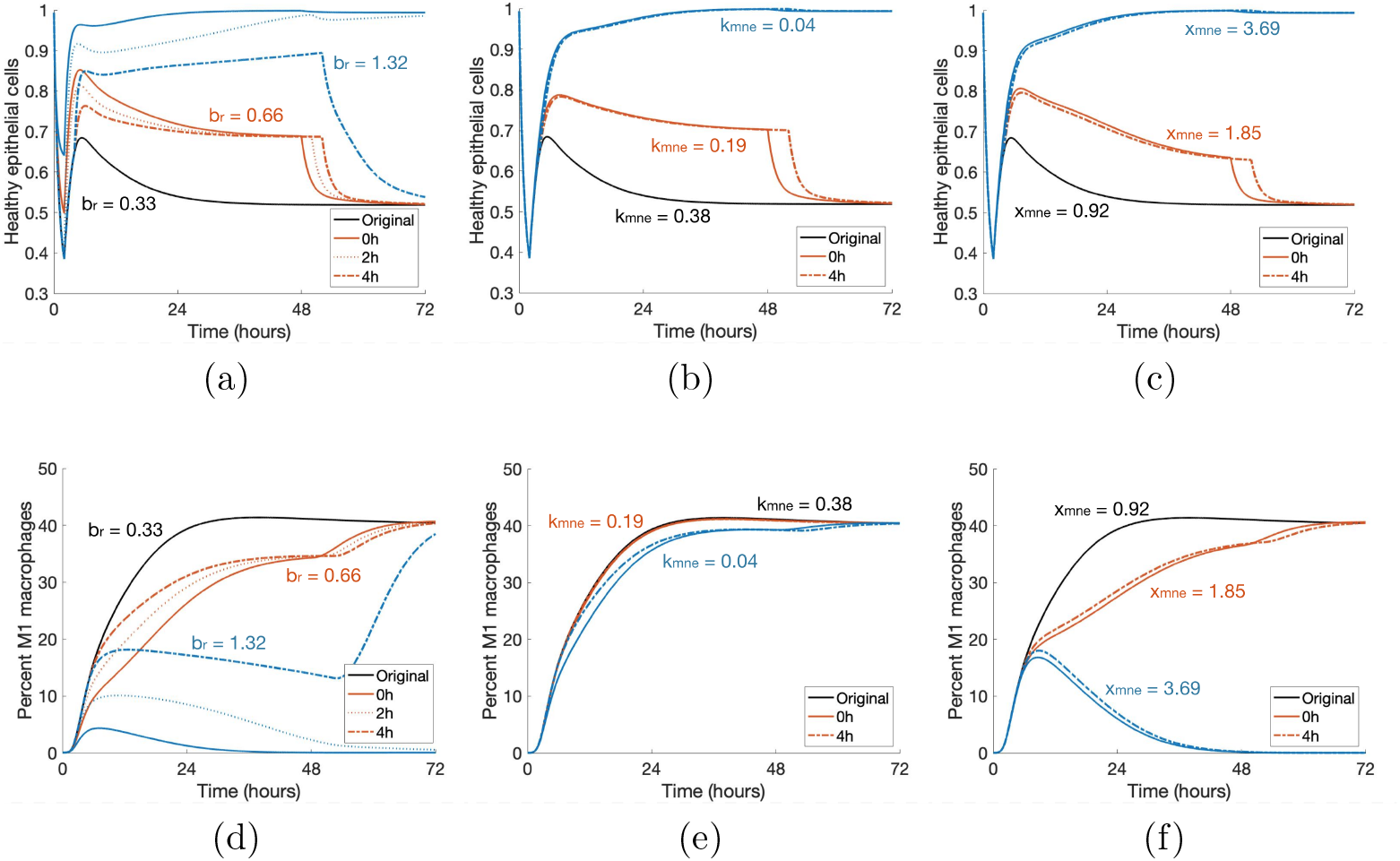
Modulating parameters based on parameter analysis improves outcome in case study. Starting with a parameter set that gives rise to an *E*_*h*_ transient that starts healthy and ends in a moderate inflammation state, we applied various treatment strategies by changing three key parameters, *b*_*r*_ (rate at which healthy epithelial cells self-repair), *k*_*mne*_ (rate of collateral damage to epithelial cells by macrophages and neutrophils), and *x*_*mne*_ (Hill-type constant which regulates the effectiveness of macrophages and neutrophils in damaging epithelial cells). Results for various changes are shown for healthy epithelial cells (a, b, c) and percent of M1 macrophages (d, e, f). Treatment was started at 0, 2, or 4 hours after the start of ventilation, denoted by solid, dotted, and dot-dashed lines, respectively, and lasted for 48 hours. The original parameter values are *b*_*r*_ = 0.33, *k*_*mne*_ = 0.38, and *x*_*mne*_ = 0.92. Black transients show the original dynamics without intervention. Orange transients represent values of each parameter that are insufficient to mediate prolonged macrophage activation. Blue transients show values that are sufficient to bring about resolution, depending on intervention time.

We intervened in a case that starts healthy and ends in moderate inflammation. Note in Fig 11, the original *E*_*h*_ transient begins recovery to healthy after the two-hour ventilation period, but by the end of the 200-hour period, is at a lower *E*_*h*_ value. This is coupled with a transient for M1 in which the pro-inflammatory phenotype increases to 40-45% and stays in this range.

Increasing *b*_*r*_ by various amounts has increasingly positive effects on long-term epithelial health. Lower values of *b*_*r*_ increase *E*_*h*_ slightly and an earlier intervention can generate a higher peak of *E*_*h*_ around five hours, but does not continue increasing at this rate regardless of intervention time. If *b*_*r*_ is increased substantially for a significant duration of treatment time, healthy epithelial cells reach the healthy steady-state after ventilation and do not decrease again. Shown in Figures 11a and 11d, doubling *b*_*r*_ to 0.66 is not enough to generate recovery, but increasing *b*_*r*_ by a factor of four to 1.32 does result in a healthy outcome. For an insufficient treatment duration and value of *b*_*r*_, levels of *E*_*h*_ will be higher until treatment ends and then decrease back to the same level as the original simulation. For a long enough treatment duration, the proportion of healthy epithelial cells will remain high even after treatment ends. For *b*_*r*_ = 0.66, the intervention time does not improve health in the long run, whereas for *b*_*r*_ = 1.32, intervention at either 0 or 2 hours is sufficient to bring about recovery while intervention at 4 hours is not.

The parameter *k*_*mne*_ has an inverse relationship with epithelial health; thus, decreasing the parameter provides better results. Decreasing *k*_*mne*_ slightly can increase the rate of recovery slightly but not enough to change the outcome to resolved. However, with a significant enough decrease of *k*_*mne*_, M1 activation peaks around hour 10 and decreases back to its original levels. The original simulation shows M1 activation leveling off at a high percentage of activation (Fig 11e). The modulated return to baseline levels is paired with a healthy outcome for epithelial cells (Fig 11b). For higher values of *k*_*mne*_, results are about the same for any intervention time 4 hours or less after the beginning of ventilation. Note in Fig 11 that the time at which intervention begins matters somewhat for changes in *b*_*r*_ but not for *k*_*mne*_. Figures 11b and 11e show that half of the original value of *k*_*mne*_ (0.38 to 0.19) is not low enough to change the outcome; multiplying by a factor of 0.1 to *k*_*mne*_ = 0.04, on the other hand, is sufficient to change the outcome to healthy.

We also increase the parameter *x*_*mne*_. Increasing this value causes the presence of macrophages and neutrophils to be less effective in damaging epithelial cells. Similarly to the other treatments, sufficient changes to *x*_*mne*_ bring about long-term recovery and the time at which intervention begins is not as important. Figures 11c and 11f show doubling *x*_*mne*_ to 1.85 is insufficient to change the outcome, and increasing *x*_*mne*_ by a factor of four to 3.69 is sufficient.

Finally, we increase *k*_*en*_. This increases the rate at which neutrophils phagocytize damaged cells, making room for new, healthy cells. Interestingly, although *k*_*en*_ is shown to be an important parameter in our analysis, even increasing the parameter by a factor of ten to 1.52 is insufficient to make any real changes in the epithelial and macrophage populations. Since there was no significant change, we do not show this treatment in Fig 11.

We also examine the results of combination therapy that could include regulation of two or three parameters. Together, changes in parameter values that would be insufficient on their own are able to regulate macrophage activation and bring epithelial cells back to a healthy state. Additionally, higher values of *b*_*r*_ and *x*_*mne*_ and lower values of *k*_*mne*_ precipitate a quicker recovery from damage. Intervention time is important for parameter values near the threshold, but not for parameter values sufficiently above or below the threshold. Intervention time may make a difference in the ending steady-state values of *E*_*h*_ or *M*1, depending on the parameters. Many combinations could be formulated; Fig 12 shows two cases in which two parameter changes were insufficient to bring about recovery individually but are sufficient when combined. The orange curves show *b*_*r*_ = 0.99 and *k*_*mne*_ = 0.19 and the blue curves show *x*_*mne*_ = 2.31 and *k*_*en*_ = 1.52, which bring about long-term recovery for all three intervention times.

**Figure 12:**
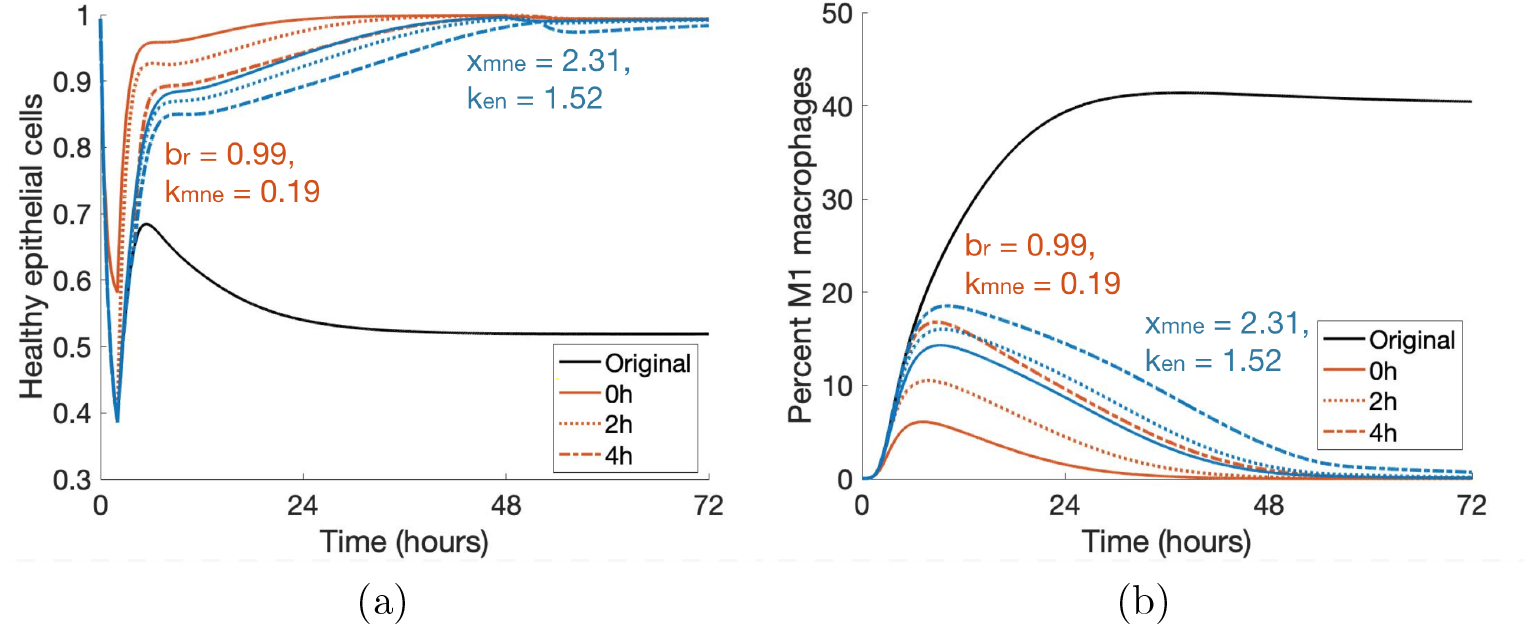
Treatment by combining parameter changes can result in a positive outcome. Changes in *b*_*r*_, *k*_*mne*_, *x*_*mne*_ and *k*_*en*_ that are insufficient on their own (Fig 11) result in a change in outcome when combined. Orange curves show a combination treatment of *b*_*r*_ = 0.99 and *k*_*mne*_ = 0.19 and blue curves show that of *x*_*mne*_ = 2.31 and *k*_*en*_ = 1.52. Duration of treatment in each case is 48 hours, and all intervention times are successful in a long-term recovery.

For other cases starting in a healthy state and ending in moderate inflammation or severe inflammation, a high enough *b*_*r*_ can bring about resolution in some cases. In general, earlier intervention times result in a faster rate of recovery, but there are varied responses to changes in *k*_*mne*_, *x*_*mne*_, and *k*_*en*_. Even for transients with similar *E*_*h*_ and *M* 1 dynamics, reactions to treatments may be different, reinforcing the uniqueness of each individual member of the virtual cohort.

## 4. Discussion

The spectrum of macrophage activation has been a recently growing field of research [10, 14, 15], and with the increase in the need for mechanical ventilation due to COVID-19, a better understanding of and treatment for VILI is of great concern. Mathematical models have studied a host of causes of lung inflammation, including bacterial and viral infections and allergic reactions. Our model combines the varied effects of macrophage activation with a more detailed epithelial subsystem to model ventilator-induced lung injury. These features help to provide a better understanding of how the components of immune response, including those associated with the different macrophage phenotypes, play a role in whether or not there is resolution after ventilator-induced damage.

We account for recruitment of circulating immune cells from the blood-stream and their contribution to the immune response using a two-compartmental model. Our model incorporates a number of factors involved in the immune response, including naive M0, pro-inflammatory M1 and anti-inflammatory M2 macrophages, three states of epithelial cells (healthy, damaged, dead), activated and unactivated neutrophils, and various mediators used to signal between cells. The model consists of 18 state equations and 67 parameters. Because of its large size and the paucity of experimental data, we used Latin hypercube sampling to find biologically meaningful parameter sets, producing a total of 23,086 acceptable parameter sets. This “virtual cohort” produces a variety of dynamics that can be generated by the model. We classified parameter sets into categories of healthy, moderate inflammation, and severe inflammation based on the percentage of healthy epithelial cells at the beginning or end of the simulation. The resulting cohort simulations are used to determine the unique characteristics and properties of the transients that are linked to outcome and to determine candidate treatments.

We utilized several methods to determine the most important parameters for model output, particularly epithelial health. Using eFAST, a sensitivity analysis method for non-linear, non-monotonic ODEs, we found parameters that, when fluctuated, caused a statistically significant difference in output than that generated by a dummy parameter. We then compared these results with more non-conventional and less computationally intensive methods. The random forest decision tree algorithm generated values denoting the importance of parameters and other predictors on epithelial health and is particularly useful for large data sets, such as the parameter sets in our virtual cohort. Additionally, significance testing determined statistically significant differences in parameters grouped by outcome.

We were able to not only include parameter values in this analysis but also other predictors later found to be important, including the M1 peak ratio and the difference between *E*_*h*_ initial condition and ending value. Three of the most important parameters were *b*_*r*_, the rate of self-repair of epithelial cells, *k*_*mne*_, the rate at which macrophages and neutrophils cause collateral damage to epithelial cells, *x*_*mne*_, the Hill-type coefficient that regulates the effectiveness of that collateral damage, and *k*_*en*_, the rate of phagocytosis of damaged epithelial cells by neutrophils. These important parameters and predictors were confirmed by at least two of the methods used.

Analysis showed that properties and parameters related to epithelial repair and M1 activation and de-activation were especially predictive of outcome. We used *b*_*r*_, *k*_*mne*_, *x*_*mne*_, and *k*_*en*_ to simulate treatments for a parameter set in the virtual cohort that started healthy and ended in a moderate inflammation disease progression. We found that modulating *b*_*r*_ is effective in most cases, and the other four can be helpful in some. The chosen case responded differently to treatments and these were paired with varied M1 activation dynamics, indicating that macrophage activation is tied to epithelial health in VILI.

Our approach of developing a virtual cohort and selecting important parameters is a first step in identifying the driving mechanisms behind VILI and how they contribute to outcomes. However, experimental data will be necessary to better understand the immune response to VILI and identify biologically realistic dynamics. Concentrations of macrophages and neutrophils, as well as a way to experimentally measure epithelial health at multiple time points would be extremely beneficial. Preliminary data is currently being collected, which can be explored in future work.

Another area of further study is determining why some virtual cases can recover with a short intervention time while others need indefinite treatment. We hypothesize that this has to do with patient-specific initial conditions and parameters but more work should be done to obtain a definite answer. This would help determine the risk of VILI for patients who undergo ventilation, since patients generally need ventilation because of a preexisting condition and do not begin ventilation in a completely healthy state. In fact, this model could be extended to include other types of injury such as a bacterial or viral infection to study the interactions between the different types of injury and how they contribute to patient outcome.

In conclusion, our model contributes to the current understanding of the immune response in the lungs, and is an important first step for VILI. Our parameter analysis using a variety of methods provides new insight into potential interventions during and after ventilation to mediate VILI. Experimental data will greatly improve our ability to suggest treatments. Furthermore, the model can be extended to include other types of injury that create the need for mechanical ventilation in the first place.

## 5. Acknowledgments

This work was supported by the National Science Foundation via award CMMI-1351162 and by the National Institutes of Health via award R21HL146250 (R.H.).

## Supplementary Material

|  |  |  |  |  |
| --- | --- | --- | --- | --- |
| A summary of the initial sates and disease progression outcomes and how they change, depending on the maximum initial amount of Ee allowed (exclusion group). |  |  |  |  |
| Initial condition criteria: |  | Ee(0)<75% | Ee(0)<50% | Ee(0)<25% |
| Total number of sets that reached steady-state: |  | 23433 | 23086 | 22217 |
|  | Healthy IC: | 15890 (438) | 15890 (438) | 15890 (428) |
|  | Health ES: | 15454 (2) | 15453 (1) | 15452 (0) |
| Moderate inflammation IC: |  | 5090 (101) | 5090 (101) | 4433 (37) |
| Moderate inflammation ES: |  | 5320 (331) | 5319 (330) | 4726 (330) |
| Severe inflammation IC: |  | 2453 (2) | 2106 (0) | 1894 (0) |
| Severe inflammation ES: |  | 2659 (208) | 2314 (208) | 2039 (145) |
| Numbers in parentheses are the number of sets that leave the state and enter the state at the end of the simulation for initial condition (IC) and ending state (ES), respectively. |  |  |  |  |

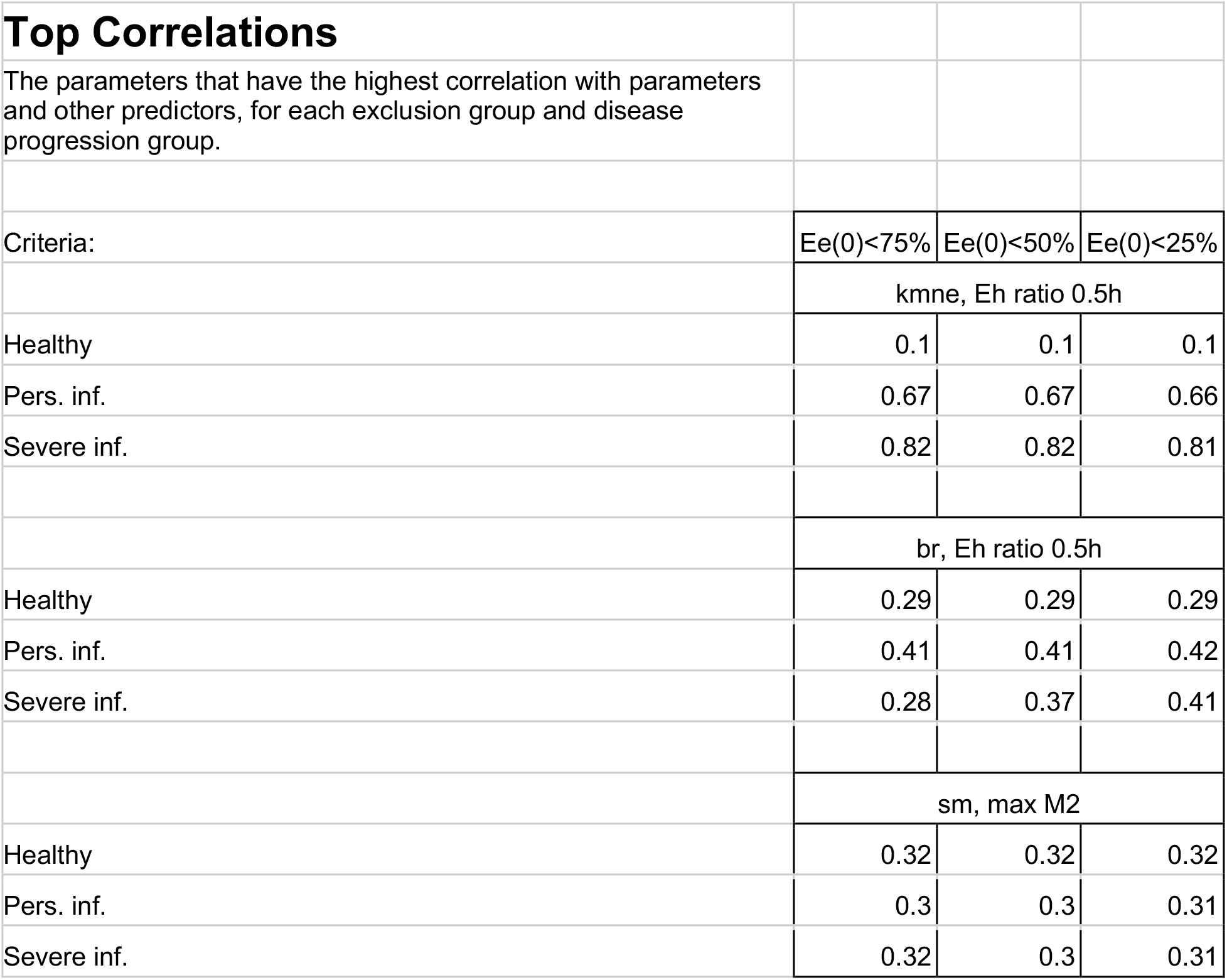

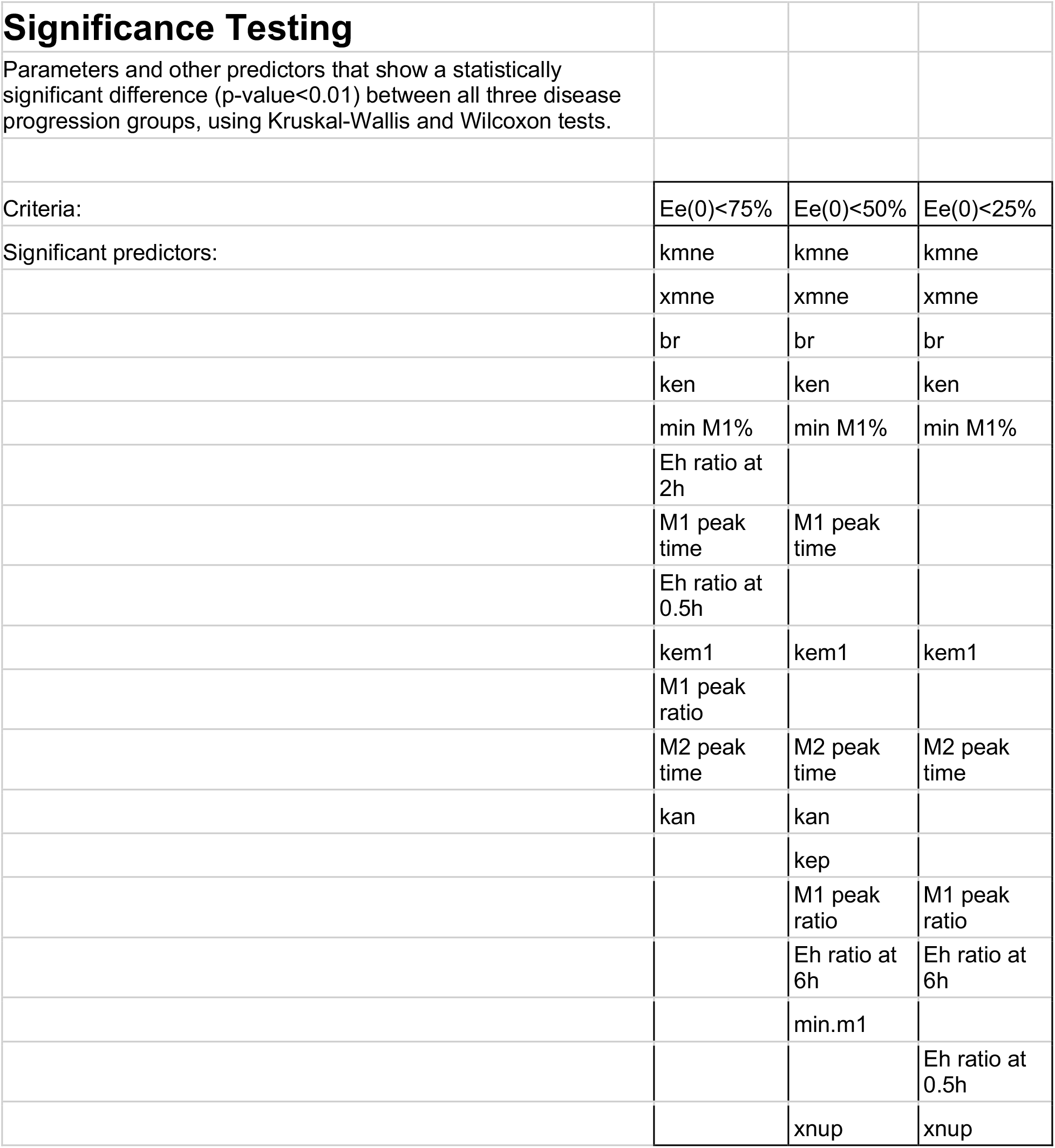

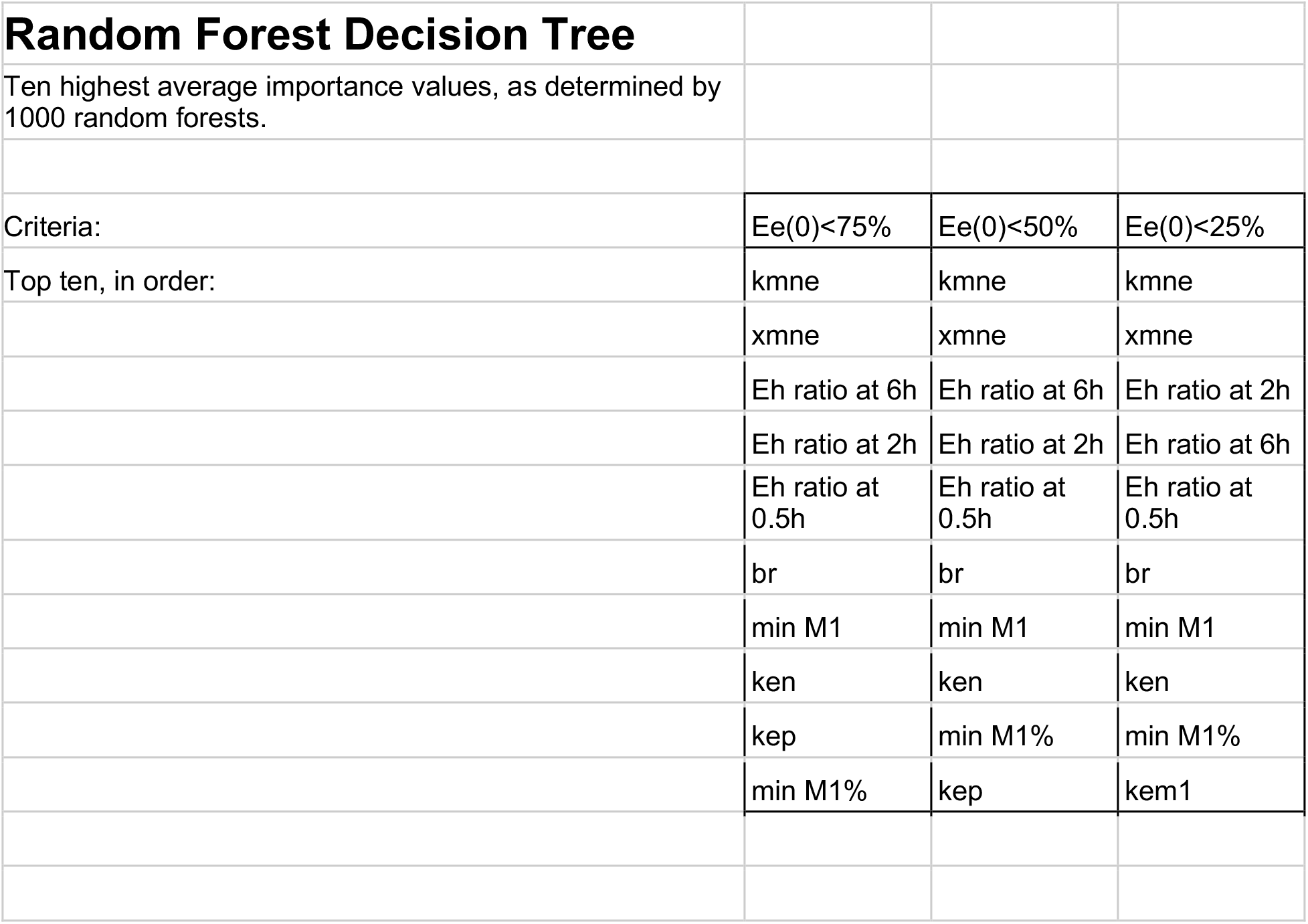

## References

[1] E. Mahase, Covid-19: most patients require mechanical ventilation in first 24 hours of critical care, BMJ 368 (2020). URL: https://www.bmj.com/content/368/bmj.m1201. doi:10.1136/bmj.m1201, publisher: British Medical Journal Publishing Group Section: News.

[2] F. J. J. Halbertsma, M. Vaneker, G. J. Scheffer, J. G. van der Hoeven, Cytokines and biotrauma in ventilator-induced lung injury: a critical review of the literature, The Netherlands Journal of Medicine 63 (2005) 382–392.

[3] A. S. Slutsky, V. M. Ranieri, Ventilator-Induced Lung Injury, New England Journal of Medicine 369 (2013) 2126–2136. URL: http://dx.doi.org/10.1056/NEJMra1208707. doi:10.1056/NEJMra1208707.

[4] M. Provinciali, M. Cardelli, F. Marchegiani, Inflammation, chronic obstructive pulmonary disease and aging, Current Opinion in Pulmonary Medicine 17 (2011) S3. doi:10.1097/01.mcp.0000410742.90463.1f.

[5] N. d. Rekeneire, R. Peila, J. Ding, L. H. Colbert, M. Visser, R. I. Shorr, S. B. Kritchevsky, L. H. Kuller, E. S. Strotmeyer, A. V. Schwartz, B. Vellas, T. B. Harris, Diabetes, Hyperglycemia, and Inflammation in Older Individuals: The Health, Aging and Body Composition study, Diabetes Care 29 (2006) 1902–1908. URL: https://care.diabetesjournals.org/content/29/8/1902. doi:10.2337/dc05-2327.

[6] C. H. Canan, N. S. Gokhale, B. Carruthers, W. P. Lafuse, L. S. Schlesinger, J. B. Torrelles, J. Turner, Characterization of Lung Inflammation and its Impact on Macrophage Function in Aging, Journal of Leukocyte Biology 96 (2014) 473–480. URL: http://www.jleukbio.org.proxy.library.vcu.edu/content/96/3/473. doi:10.1189/jlb.4A0214-093RR.

[7] Y. Feng, Y. Amoateng-Adjepong, D. Kaufman, C. Gheorghe, C. A. Manthous, Age, duration of mechanical ventilation, and outcomes of patients who are critically ill, Chest 136 (2009) 759–764.

[8] Z. Wu, J. M. McGoogan, Characteristics of and Important Lessons From the Coronavirus Disease 2019 (COVID-19) Outbreak in China: Summary of a Report of 72 314 Cases From the Chinese Center for Disease Control and Prevention, JAMA 323 (2020) 1239–1242. URL: https://jamanetwork.com/journals/jama/fullarticle/2762130. doi:10.1001/jama.2020.2648, publisher: American Medical Association.

[9] G. Bruno, S. Perelli, C. Fabrizio, G. B. Buccoliero, Short-term outcomes in individuals aged 75 or older with severe coronavirus disease (COVID-19): First observations from an Infectious Diseases Unit in Southern Italy, The Journal of Infection (2020). URL: https://www.ncbi.nlm.nih.gov/pmc/articles/PMC7224683/. doi:10.1016/j.jinf.2020.05.024.

[10] M. Torres, J. Wang, P. J. Yannie, S. Ghosh, R. A. Segal, A. M. Reynolds, Identifying important parameters in the inflammatory process with a mathematical model of immune cell influx and macrophage polarization, PLOS Computational Biology 15 (2019) e1007172. URL: https://journals.plos.org/ploscompbiol/article?id=10.1371/journal.pcbi.1007172 doi:10.1371/journal.pcbi.1007172.

[11] M. A. Matthay, L. Robriquet, X. Fang, Alveolar Epithelium, Proceedings of the American Thoracic Society 2 (2005) 206–213. URL: https://www.atsjournals.org/doi/full/10.1513/pats.200501-009AC. doi:10.1513/pats.200501-009AC.

[12] R. J. Mason, Biology of alveolar type II cells, Respirology 11 (2006) S12–S15. URL: https://onlinelibrary.wiley.com/doi/abs/10.1111/j.1440-1843.2006.00800.x. doi:10.1111/j.1440-1843.2006.00800.x.

[13] L. B. Ware, M. A. Matthay, The Acute Respiratory Distress Syndrome, New England Journal of Medicine 342 (2000) 1334–1349. URL: https://doi.org/10.1056/NEJM200005043421806. doi:10.1056/NEJM200005043421806.

[14] N. R. Aggarwal, L. S. King, F. R. D’Alessio, Diverse Macrophage Populations Mediate Acute Lung Inflammation and Resolution, American Journal of Physiology-Lung Cellular and Molecular Physiology 306 (2014) L709–L725. URL: http://www.physiology.org/doi/abs/10.1152/ajplung.00341.2013. doi:10.1152/ajplung.00341.2013.

[15] D. M. Mosser, J. P. Edwards, Exploring the Full Spectrum of Macrophage Activation, Nature Reviews Immunology 8 (2008) 958–969. URL: http://www.nature.com.proxy.library.vcu.edu/nri/journal/v8/n12/full/nri2448.ht doi:10.1038/nri2448.

[16] N. Wang, H. Liang, K. Zen, Molecular Mechanisms That Influence the Macrophage M1–M2 Polarization Balance, Frontiers in Immunology 5 (2014). URL: http://www.ncbi.nlm.nih.gov/pmc/articles/PMC4246889/. doi:10.3389/fimmu.2014.00614.

[17] E. Kolaczkowska, P. Kubes, Neutrophil Recruitment and Function in Health and Inflammation, Nature Reviews Immunology 13 (2013) 159–175. URL: https://www.nature.com/articles/nri3399. doi:10.1038/nri3399.

[18] J. Grommes, O. Soehnlein, Contribution of Neutrophils to Acute Lung Injury, Molecular Medicine 17 (2011) 293–307. URL: https://www.ncbi.nlm.nih.gov/pmc/articles/PMC3060975/. doi:10.2119/molmed.2010.00138.

[19] E. J. Naylor, D. Bakstad, M. Biffen, B. Thong, P. Calverley, S. Scott, C. A. Hart, R. J. Moots, S. W. Edwards, Haemophilus Influenzae Induces Neutrophil Necrosis, American Journal of Respiratory Cell and Molecular Biology 37 (2007) 135–143. URL: https://www.atsjournals.org/doi/full/10.1165/rcmb.2006-0375OC. doi:10.1165/rcmb.2006-0375OC.

[20] E. Linehan, D. Fitzgerald, Ageing and the Immune System: Focus on Macrophages, European Journal of Microbiology and Immunology 5 (2015) 14–24. URL: http://akademiai.com/doi/abs/10.1556/EuJMI-D-14-00035. doi:10.1556/EuJMI-D-14-00035.

[21] S. Mahbub, C. R. Deburghgraeve, E. J. Kovacs, Advanced Age Impairs Macrophage Polarization, Journal of Interferon & Cytokine Research 32 (2011) 18–26. URL: http://online.liebertpub.com/doi/abs/10.1089/jir.2011.0058. doi:10.1089/jir.2011.0058.

[22] S. Schirm, P. Ahnert, S. Wienhold, H. Mueller-Redetzky, G. Nouailles-Kursar, M. Loeffler, M. Witzenrath, M. Scholz, A Biomathematical Model of Pneumococcal Lung Infection and Antibiotic Treatment in Mice, PLOS ONE 11 (2016) e0156047. URL: http://journals.plos.org/plosone/article?id=10.1371/journal.pone.0156047. doi:10.1371/journal.pone.0156047.

[23] E. Mochan, D. Swigon, G. B. Ermentrout, S. Lukens, G. Clermont, A Mathematical Model of Intrahost Pneumococcal Pneumonia Infection Dynamics in Murine Strains, Journal of Theoretical Biology 353 (2014) 44–54. URL: http://www.sciencedirect.com/science/article/pii/S0022519314000964. doi:10.1016/j.jtbi.2014.02.021.

[24] A. M. Smith, J. A. McCullers, F. R. Adler, Mathematical Model of a Three-Stage Innate Immune Response to a Pneumococcal Lung Infection, Journal of Theoretical Biology 276 (2011) 106–116. URL: http://www.sciencedirect.com/science/article/pii/S0022519311000786. doi:10.1016/j.jtbi.2011.01.052.

[25] J. Day, A. Friedman, L. S. Schlesinger, Modeling the Immune Rheostat of Macrophages in the Lung in Response to Infection, Proceedings of the National Academy of Sciences 106 (2009) 11246–11251. URL: http://www.pnas.org.proxy.library.vcu.edu/content/106/27/11246. doi:10.1073/pnas.0904846106.

[26] K. Raman, A. G. Bhat, N. Chandra, A Systems Perspective of Host–Pathogen Interactions: Predicting Disease Outcome in Tuberculosis, Molecular BioSystems 6 (2010) 516–530. URL: http://pubs.rsc.org/en/content/articlelanding/2010/mb/b912129c. doi:10.1039/B912129C.

[27] J. L. Segovia-Juarez, S. Ganguli, D. Kirschner, Identifying Control Mechanisms of Granuloma Formation During M. tuberculosis Infection Using an Agent-Based Model, Journal of Theoretical Biology 231 (2004) 357–376. URL: http://www.sciencedirect.com/science/article/pii/S0022519304003212. doi:10.1016/j.jtbi.2004.06.031.

[28] H. Manchanda, N. Seidel, A. Krumbholz, A. Sauerbrei, M. Schmidtke, R. Guthke, Within-Host Influenza Dynamics: A Small-Scale Mathematical Modeling Approach, Biosystems 118 (2014) 51–59. URL: http://www.sciencedirect.com/science/article/pii/S0303264714000331. doi:10.1016/j.biosystems.2014.02.004.

[29] C. S. Anderson, M. L. DeDiego, D. J. Topham, J. Thakar, Boolean Modeling of Cellular and Molecular Pathways Involved in Influenza Infection, Computational and Mathematical Methods in Medicine 2016 (2016). URL: https://www.ncbi.nlm.nih.gov/pmc/articles/PMC4769743/. doi:10.1155/2016/7686081.

[30] B. Hancioglu, D. Swigon, G. Clermont, A Dynamical Model of Human Immune Response to Influenza A Virus Infection, Journal of Theoretical Biology 246 (2007) 70–86. URL: http://www.sciencedirect.com/science/article/pii/S0022519306005820. doi:10.1016/j.jtbi.2006.12.015.

[31] B. N. Brown, I. M. Price, F. R. Toapanta, D. R. DeAlmeida, C. A. Wiley, T. M. Ross, T. D. Oury, Y. Vodovotz, An Agent-Based Model of Inflammation and Fibrosis Following Particulate Exposure in the Lung, Mathematical Biosciences 231 (2011) 186–196. URL: http://www.sciencedirect.com/science/article/pii/S0025556411000356. doi:10.1016/j.mbs.2011.03.005.

[32] I. L. Chernyavsky, H. Croisier, L. A. C. Chapman, L. S. Kimpton, J. E. Hiorns, B. S. Brook, O. E. Jensen, C. K. Billington, P. Hall, S. R. Johnson, The Role of Inflammation Resolution Speed in Airway Smooth Muscle Mass Accumulation in Asthma: Insight from a Theoretical Model, PLOS ONE 9 (2014) e90162. URL: http://journals.plos.org/plosone/article?id=10.1371/journal.pone.0090162. doi:10.1371/journal.pone.0090162.

[33] A. Golov, S. Simakov, Y. N. Soe, R. Pryamonosov, O. Mynbaev, A. Kholodov, Multiscale CT-Based Computational Modeling of Alveolar Gas Exchange during Artificial Lung Ventilation, Cluster (Biot) and Periodic (Cheyne-Stokes) Breathings and Bronchial Asthma Attack, Computation 5 (2017) 11. URL: http://www.mdpi.com/2079-3197/5/1/11. doi:10.3390/computation5010011.

[34] J. J. Pothen, M. E. Poynter, J. H. T. Bates, A Computational Model of Unresolved Allergic Inflammation in Chronic Asthma, American Journal of Physiology - Lung Cellular and Molecular Physiology 308 (2015) L384–L390. URL: http://ajplung.physiology.org.proxy.library.vcu.edu/content/308/4/L384. doi:10.1152/ajplung.00268.2014.

[35] P. Aghasafari, I. Bin M. Ibrahim, R. Pidaparti, Strain-induced inflammation in pulmonary alveolar tissue due to mechanical ventilation, Biomechanics and Modeling in Mechanobiology 16 (2017) 1103–1118. URL: https://doi.org/10.1007/s10237-017-0879-5. doi:10.1007/s10237-017-0879-5.

[36] J. H. T. Bates, C. G. Irvin, Time dependence of recruitment and derecruitment in the lung: a theoretical model, Journal of Applied Physiology 93 (2002) 705–713. URL: https://journals.physiology.org/doi/full/10.1152/japplphysiol.01274.2001. doi:10.1152/japplphysiol.01274.2001, publisher: American Physiological Society.

[37] K. L. Hamlington, B. J. Smith, G. B. Allen, J. H. T. Bates, Predicting ventilator-induced lung injury using a lung injury cost function, Journal of Applied Physiology 121 (2016) 106–114. URL: https://journals.physiology.org/doi/full/10.1152/japplphysiol.00096.2016. doi:10.1152/japplphysiol.00096.2016, publisher: American Physiological Society.

[38] K. G. Hickling, The Pressure–Volume Curve Is Greatly Modified by Recruitment, American Journal of Respiratory and Critical Care Medicine 158 (1998) 194–202. URL: http://www.atsjournals.org.proxy.library.vcu.edu/doi/full/10.1164/ajrccm.158.1 doi:10.1164/ajrccm.158.1.9708049.

[39] J. Kim, R. L. Heise, A. M. Reynolds, R. M. Pidaparti, Quantification of Age-Related Lung Tissue Mechanics under Mechanical Ventilation, Medical Sciences 5 (2017) 21. URL: https://www.mdpi.com/2076-3271/5/4/21. doi:10.3390/medsci5040021, number: 4 Publisher: Multidisciplinary Digital Publishing Institute.

[40] J. J. Marini, P. S. Crooke, J. D. Truwit, Determinants and Limits of Pressure-Preset Ventilation: a Mathematical Model of Pressure Control, Journal of Applied Physiology 67 (1989) 1081–1092. URL: http://jap.physiology.org.proxy.library.vcu.edu/content/67/3/1081.

[41] C. B. Massa, G. B. Allen, J. H. T. Bates, Modeling the dynamics of recruitment and derecruitment in mice with acute lung injury, Journal of Applied Physiology 105 (2008) 1813–1821. URL: https://journals.physiology.org/doi/full/10.1152/japplphysiol.90806.2008. doi:10.1152/japplphysiol.90806.2008, publisher: American Physiological Society.

[42] R. M. Pidaparti, M. Burnette, R. L. Heise, A. Reynolds, Analysis for Stress Environment in the Alveolar Sac Model, Journal of biomedical science and engineering 6 (2013) 901–907. URL: http://www.ncbi.nlm.nih.gov/pmc/articles/PMC4057278/. doi:10.4236/jbise.2013.69110.

[43] A. Reynolds, G. Bard Ermentrout, G. Clermont, A Mathematical Model of Pulmonary Gas Exchange Under Inflammatory Stress, Journal of Theoretical Biology 264 (2010) 161–173. URL: http://www.sciencedirect.com/science/article/pii/S0022519310000159. doi:10.1016/j.jtbi.2010.01.011.

[44] D. A. Braun, M. Fribourg, S. C. Sealfon, Cytokine Response Is Determined by Duration of Receptor and Signal Transducers and Activators of Transcription 3 (STAT3) Activation, Journal of Biological Chemistry 288 (2013) 2986–2993. URL: http://www.jbc.org/content/288/5/2986. doi:10.1074/jbc.M112.386573.

[45] S. Maiti, W. Dai, R. C. Alaniz, J. Hahn, A. Jayaraman, Mathematical Modeling of Pro- and Anti-Inflammatory Signaling in Macrophages, Processes 3 (2014) 1–18. URL: http://www.mdpi.com/2227-9717/3/1/1. doi:10.3390/pr3010001.

[46] L. M. Crosby, C. M. Waters, Epithelial Repair Mechanisms in the Lung, American Journal of Physiology-Lung Cellular and Molecular Physiology 298 (2010) L715–L731. URL: http://www.physiology.org/doi/abs/10.1152/ajplung.00361.2009. doi:10.1152/ajplung.00361.2009.

[47] A. Gardner, L. A. Borthwick, A. J. Fisher, Lung Epithelial Wound Healing in Health and Disease, Expert Review of Respiratory Medicine 4 (2010) 647–660. URL: https://doi.org/10.1586/ers.10.62. doi:10.1586/ers.10.62.

[48] S. Herold, K. Mayer, J. Lohmeyer, Acute Lung Injury: How Macrophages Orchestrate Resolution of Inflammation and Tissue Repair, Frontiers in Immunology 2 (2011). URL: https://www.ncbi.nlm.nih.gov/pmc/articles/PMC3342347/. doi:10.3389/fimmu.2011.00065.

[49] B. Ermentrout, Simulating, Analyzing, and Animating Dynamical Systems, Software, Environments and Tools, Society for Industrial and Applied Mathematics, 2002. URL: http://epubs.siam.org/doi/book/10.1137/1.9780898718195. doi:10.1137/1.9780898718195.

[50] C. Nathan, Neutrophils and Immunity: Challenges and Opportunities, Nature Reviews Immunology 6 (2006) 173–182. URL: https://www.nature.com/articles/nri1785. doi:10.1038/nri1785.

[51] S. Gordon, Alternative Activation of Macrophages, Nature Reviews Immunology 3 (2003) 23–35. URL: http://www.nature.com/articles/nri978. doi:10.1038/nri978.

[52] C. T. Robb, K. H. Regan, D. A. Dorward, A. G. Rossi, Key Mechanisms Governing Resolution of Lung Inflammation, Seminars in Immunopathology 38 (2016) 425–448. URL: https://link-springer-com.proxy.library.vcu.edu/article/10.1007/s00281-016-056 doi:10.1007/s00281-016-0560-6.

[53] V. Kumar, A. Sharma, Neutrophils: Cinderella of Innate Immune System, International Immunopharmacology 10 (2010) 1325–1334. URL: http://www.sciencedirect.com/science/article/pii/S1567576910002663. doi:10.1016/j.intimp.2010.08.012.

[54] L. K. Johnston, C. R. Rims, S. E. Gill, J. K. McGuire, A. M. Manicone, Pulmonary Macrophage Subpopulations in the Induction and Resolution of Acute Lung Injury, American Journal of Respiratory Cell and Molecular Biology (2012). URL: http://www.atsjournals.org/doi/abs/10.1165/rcmb.2012-0090OC. doi:10.1165/rcmb.2012-0090OC.

[55] C. Summers, S. M. Rankin, A. M. Condliffe, N. Singh, A. M. Peters, E. R. Chilvers, Neutrophil Kinetics in Health and Disease, Trends in Immunology 31 (2010) 318–324. URL: http://www.sciencedirect.com/science/article/pii/S147149061000075X. doi:10.1016/j.it.2010.05.006.

[56] S. M. Opal, V. A. DePalo, Anti-inflammatory cytokines, Chest 117 (2000) 1162–1172.

[57] N. E. Vlahakis, M. A. Schroeder, A. H. Limper, R. D. Hubmayr, Stretch Induces Cytokine Release by Alveolar Epithelial Cells in Vitro, American Journal of Physiology-Lung Cellular and Molecular Physiology 277 (1999) L167–L173. URL: http://www.physiology.org/doi/abs/10.1152/ajplung.1999.277.1.L167. doi:10.1152/ajplung.1999.277.1.L167.

[58] M. Heusinkveld, P. J. d. V. v. Steenwijk, R. Goedemans, T. H. Ramwadhdoebe, A. Gorter, M. J. P. Welters, T. v. Hall, S. H. v. d. Burg, M2 Macrophages Induced by Prostaglandin E2 and IL-6 from Cervical Carcinoma Are Switched to Activated M1 Macrophages by CD4+ Th1 Cells, The Journal of Immunology 187 (2011) 1157–1165. URL: http://www.jimmunol.org/content/187/3/1157. doi:10.4049/jimmunol.1100889.

[59] O. Soehnlein, L. Lindbom, Phagocyte partnership during the onset and resolution of inflammation, Nature Reviews Immunology 10 (2010) 427–439. URL: https://www.nature.com/articles/nri2779. doi:10.1038/nri2779.

[60] M. D. McKay, R. J. Beckman, W. J. Conover, Comparison of Three Methods for Selecting Values of Input Variables in the Analysis of Output from a Computer Code, Technometrics 21 (1979) 239–245. doi:10.1080/00401706.1979.10489755.

[61] S. Marino, I. B. Hogue, C. J. Ray, D. E. Kirschner, A methodology for performing global uncertainty and sensitivity analysis in systems biology, Journal of Theoretical Biology 254 (2008) 178–196. URL: http://www.sciencedirect.com/science/article/pii/S0022519308001896. doi:10.1016/j.jtbi.2008.04.011.

[62] D. E. Kirschner, Matlab Functions for PRCC and eFAST, 2008. URL: http://malthus.micro.med.umich.edu/lab/usadata/.

[63] L. S. Smith, S. A. Gharib, C. W. Frevert, T. R. Martin, Effects of Age on the Synergistic Interactions between Lipopolysaccharide and Mechanical Ventilation in Mice, American Journal of Respiratory Cell and Molecular Biology 43 (2010) 475–486. URL: https://www.atsjournals.org/doi/full/10.1165/rcmb.2009-0039OC. doi:10.1165/rcmb.2009-0039OC, publisher: American Thoracic Society - AJRCMB.

[64] E. K. Wolthuis, A. P. Vlaar, G. Choi, J. J. Roelofs, N. P. Juffermans, M. J. Schultz, Mechanical ventilation using non-injurious ventilation settings causes lung injury in the absence of pre-existing lung injury in healthy mice, Critical Care 13 (2009) R1. URL: https://doi.org/10.1186/cc7688. doi:10.1186/cc7688.

[65] A. Saltelli, S. Tarantola, F. Campolongo, M. Ratto, Sensitivity Analysis in Practice: A Guide to Assessing Scientific Models, John Wiley & Sons, 2004. Google-Books-ID: NsAVmohPNpQC.

[66] A. Saltelli, R. Bolado, An alternative way to compute Fourier amplitude sensitivity test (FAST), Computational Statistics & Data Analysis 26 (1998) 445–460. URL: http://www.sciencedirect.com/science/article/pii/S0167947397000431. doi:10.1016/S0167-9473(97)00043-1.

[67] A. Saltelli, S. Tarantola, K. P.-S. Chan, A Quantitative Model-Independent Method for Global Sensitivity Analysis of Model Output, Technometrics 41 (1999) 39–56. URL: https://www.tandfonline.com/doi/abs/10.1080/00401706.1999.10485594. doi:10.1080/00401706.1999.10485594.

[68] R. I. Cukier, C. M. Fortuin, K. E. Shuler, A. G. Petschek, J. H. Schaibly, Study of the sensitivity of coupled reaction systems to uncertainties in rate coefficients. I Theory, The Journal of Chemical Physics 59 (1973) 3873–3878. URL: https://aip.scitation.org/doi/abs/10.1063/1.1680571. doi:10.1063/1.1680571.

[69] J. H. Schaibly, K. E. Shuler, Study of the sensitivity of coupled reaction systems to uncertainties in rate coefficients. II Applications, The Journal of Chemical Physics 59 (1973) 3879–3888. URL: https://aip.scitation.org/doi/abs/10.1063/1.1680572. doi:10.1063/1.1680572.

[70] D. C. Collins, R. Avissar, An Evaluation with the Fourier Amplitude Sensitivity Test (FAST) of Which Land-Surface Parameters Are of Greatest Importance in Atmospheric Modeling, Journal of Climate 7 (1994) 681–703.

[71] J. Le, Decision Trees in R, 2018. URL: https://www.datacamp.com/community/tutorials/decision-trees-R.

[72] A. Liaw, M. Wiener, Classification and Regression by randomForest, 2002.

[73] P. E. McKight, J. Najab, Kruskal-Wallis Test, in: The Corsini Encyclopedia of Psychology, American Cancer Society, 2010, pp. 1–1. URL: https://onlinelibrary.wiley.com/doi/abs/10.1002/9780470479216.corpsy0491. doi:10.1002/9780470479216.corpsy0491.

[74] Y. Benjamini, Y. Hochberg, Controlling the false discovery rate: a practical and powerful approach to multiple testing, Journal of the Royal statistical society: series B (Methodological) 57 (1995) 289–300.

