## Supplemental Material for "Mathematical modeling of ventilator-induced lung inflammation" (from another version of this preprint: version 1, 2020-06-04)

### Supporting Information

#### Understanding the Role of Macrophages in Lung Inflammation Through Mathematical Modeling

Sarah B. Minucci[a], Rebecca L. Heise[b, c], Michael S. Valentine[b], Franck J. Kamga Gninzeko[b], Angela M. Reynolds*[a, c]
[a]Virginia Commonwealth University, Department of Mathematics & Applied Mathematics, Richmond, Virginia
[b]Virginia Commonwealth University, Department of Biomedical Engineering, Richmond, Virginia
[c]Victoria Johnson Center, Virginia Commonwealth University, Department of Mathematics & Applied Mathematics, Richmond, Virginia

**Table S1:** List of antibodies, clones, manufacturers, and concentrations for flow cytometric analysis.

| Marker | Alveolar Macrophages | Interstitial Macrophages | Clone | Isotype | Fluorochrome Conjugate | Dilution Factor |
|---|---|---|---|---|---|---|
| CD45 | + | + | 30-F11 | Rat IgG2b,κ | BB515 | **1:100** |
| CD11b | - | + | M1/70 | Rat IgG2b,κ | BUV395 | **1:100** |
| CD11c | + | + | HL3 | Armenian Hamster IgG1, λ2 | BV786 | **1:50** |
| CD24 | - | - | M1/69 | Rat IgG2b,κ | AF700 | **1:300** |
| CD64 | + | + | X54-5/7.1 | Mouse IgG1, κ | PE | **1:200** |
| MHCII | +/- | + | M5/114.15.2 | Rat IgG2b,κ | PerCP-Cy5.5 | **1:300** |
| CD80 (M1) | +/- | +/- | 16-10A1 | Armenian Hamster IgG1, λ2 | BV421 | **1:200** |
| CD206 (M2) | +/- | +/- | MR5D3 | Rat IgG2a | AF647 | **1:200** |
